# Induction of Interferon-Stimulated Genes and Cellular Stress Pathways by Morpholinos

**DOI:** 10.1101/479188

**Authors:** Jason K.H. Lai, Kristina Gagalova, Didier Y.R. Stainier

## Abstract

The phenotypes caused by morpholino-mediated interference of gene function in zebrafish are often not observed in the corresponding mutant(s). We took advantage of the availability of a relatively large collection of transcriptomic datasets to identify common signatures that characterize morpholino-injected animals (morphants). In addition to the previously reported activation of *tp53* expression, we observed increased expression of the interferon-stimulated genes (ISGs), *isg15* and *isg20*, an extrinsic cell death pathway gene, *casp8*, and other cellular stress response genes including *phlda3*, *mdm2* and *gadd45aa*. Studies involving segmentation stage embryos were more likely to show upregulation of these genes. We also found that these genes could be upregulated by an increasing dose of *egfl7* morpholino, or high doses of standard control morpholino. Thus, these data suggest that morpholinos can stimulate the innate immune response in zebrafish, as was recently reported in *Xenopus*.

## Introduction

Morpholinos are anti-sense reagents capable of knocking down gene function by inhibiting post-transcriptional regulation of RNAs [1]. They are synthetic nucleic acid analogs which can heterodimerize with complementary RNA molecules to inhibit their processing. Morpholinos inhibit mRNAs by blocking their splicing or translation [2–3], and they can inhibit miRNA function as well [4].

The ease of morpholino use in zebrafish and its effectiveness in gene knockdown enabled scientists to transiently inhibit gene function to study various biological processes [2]. Initial testing of morpholinos indicated that morphants could largely phenocopy known mutants [2, 5]. However, the examples were limited, and restricted to genes with well characterized mutant phenotype(s). With the recent advent of gene editing, scientists have now been able to test more extensively whether reported morphant phenotypes are similar to those observed in mutants, and they have found substantial discrepancy [6]. For example, a study by Kok et al. generated mutations in more than 20 genes and observed that most mutants appear to develop normally while the corresponding morphants for ten of these genes display a developmental phenotype, suggesting additional effects introduced by morpholinos [7]. This idea was supported by the observation that the hydrocephaly phenotype in *megamind* morphants could be reproduced by injecting a *megamind* morpholino into animals devoid of the *megamind* locus, albeit at a very high dose (20 ng) [7]. In addition, Rossi, Kontarakis et al. proposed that genetic compensation in mutants but not in morphants could explain their phenotypic discrepancy [8].

One well characterized morpholino off-target effect is the activation of the p53 signaling pathway [9]. Additionally, a recent study in *Xenopus* found that morpholinos could activate the innate immune pathway, as well as induce off-target mis-splicing of pre-mRNA transcripts [10]. These studies raise questions about the effects of morpholinos in a cell, and the need to fully understand them for future applications. In this study, we sought to further characterize the effects of morpholinos by studying various microarray gene expression datasets from morpholino studies in zebrafish obtained from the Gene Expression Omnibus (GEO) database. We found that the expression of interferon-stimulated genes, ISGs (*isg15* and *isg20*), and other cellular stress pathway genes (*tp53*, *phlda3, casp8, gadd45aa* and *mdm2*) was upregulated in several morpholino studies, especially in segmentation stage embryos. Furthermore, these genes were upregulated in response to increasing doses of an *egfl7* morpholino, or high doses of a standard control morpholino. In contrast, a *vegfaa* morpholino did not induce the expression of these genes. Our study suggests that morpholinos can induce the innate immune response and furthers our understanding of morpholino effects in developing zebrafish embryos.

## Materials and Methods

### Microarray data collection and curation

Microarray gene expression arrays of various zebrafish morpholino studies were obtained from the GEO database. As of the 19th of March 2015, keyword search for “zebrafish morpholino” or “zebrafish knockdown” yielded 63 unique *Danio rerio* data series (GSE). The embedded metadata and raw intensities for all studies were retrieved with the R package GEOquery [11]. We manually curated all datasets utilizing GPL1319, a platform manufactured by Affymetrix, to annotate the publication, morpholino sequence, morpholino dose, sample source and developmental stage. Datasets from Agilent arrays were set aside for meta-analysis. All annotated information can be found in the File S1 – Section 1.

### Affymetrix microarray data normalization and processing

8 GSE data series were rejected because they were not morpholino studies, were a duplicate, or were not linked to a publication. The remaining data series were pooled together for normalization. The raw data were read with the ReadAffy function from affy package in R [12], followed by the implementation of Single Channel Array Normalization (SCAN) [13] to normalize the pooled dataset. Default parameters were used. Integrating microarray datasets from various studies will inevitably introduce batch effects. Thus, we corrected the normalized data for batch effects using ComBat [14], and defined the unique GSE ID as the primary source of batch effects.

The data were then filtered by removing control probes as well as averaging all probesets that interrogate the same gene. We also filtered genes that had little variation in gene expression across all pooled samples using the varFilter from geneFilter package. These data were used to generate the Principal Component Analysis (PCA) plots.

Finally, we grouped genes that show high correlation in the microarray dataset using a 0.95 score cutoff. This step was carried out with the findCorrelation from caret R package [15]. Details and quality control plots can be found in the File S1 – Section 2.

### Feature Selection

We tested various models with built-in feature selection on our dataset, and we also tested the final predictive accuracy. These models include: “glmnet”, “pam”, “gaussprLinear”, “svmLinear”, “svmPoly”, “svmRadial”, “nb”, “rf”, “knn” and “J48”. We optimized the parameters for each method through the train function of caret R package [15], using Leave One Out Cross Validation (LOOCV) and the accuracy as the readout score. The models were further tested against scrambled-labels datasets. Relevant genes were ranked with t-test statistic as the linear models performed best. We analyzed this set of genes by Gene Set Enrichment Analysis (GSEA) with GOrilla [16] as well as pathway analysis with Ingenuity Pathway Analysis (Qiagen).

The features were shortlisted through recursive feature elimination and backward selection with rfe caret function. The models were evaluated with 4-folds cross-validation, iterated 100 times. The model accuracy increases logarithmically with the number of features. To avoid overfitting the model, we selected the number of features at the point of diminishing returns, which is 5% below the maximum accuracy score.

### Two-way ANOVA analysis

We conducted a standard two-way ANOVA analysis stratifying for developmental stage and treatment (i.e. morphants and controls). Both variables were converted into categorical variables. To determine developmental stage, we followed ZFIN’s guidelines.

### Agilent microarray processing and analysis

A total of 34 microarray studies were done on a variety of Agilent microarray platforms. Among these studies, 12 were rejected due to the same reasons mentioned for the rejected Affymetrix studies. A further 5 raw datasets were not usable for a standard pipeline because they were uploaded to GEO in a wrong file format or were not readable. 1 other dataset could not be analyzed as the experimental design is insufficient to run statistical tests. The R package limma [17] was used to process the raw intensity files as well as downstream analyses.

Briefly, the raw files were read and compiled into a standard “eset” object for a standard normalization pipeline and statistical analyses [18]. Both dual-and single-channel arrays were first background corrected using the “normexp” method [19], followed by within-array normalization by the “loess” method to correct for dye biases in dual-channel arrays [18]. Finally, between-array normalization was performed for both microarray configurations by the “quantile” method [20]. Control probes were then removed prior to applying the empirical Bayesian statistical analyses [21]. We extracted the average log_2_-transformed fold-change (logFC) and standard errors for all probes interrogating *isg15*, *isg20*, *casp8*, *mdm2*, *gadd45aa* and *tp53*. As some genes were interrogated by multiple probes, we took the probe that detected the maximum absolute logFC.

### Ethics Statement

All zebrafish husbandry was performed under standard conditions in accordance with institutional (MPG) and national ethical and animal welfare guidelines.

### Zebrafish

Transgenic line used in this study:

*Tg(kdrl:EGFP)^s843Tg^* [22]

### Zebrafish micro-injection and Real-time PCR (qPCR) assay

The three morpholinos used in this study are *egfl7* (5’-CAGGTGTGTCTGACAGCAGAAAGAG-3’) [23], *vegfaa* (5’-GTATCAAATAAACAACCAAGTTCAT-3’) [24] and standard control morpholino (5’-CCTCTTACCTCAGTTACAATTTATA-3’) (Gene Tools). Each morpholino was injected into 1-cell stage *Tg(kdrl:EGFP)^S843Tg^* zebrafish embryos (1 nL volume). The morpholino doses used are as follows: *egfl7* - 0.0625, 0.125, 0.25, 0.5, 1 and 2 ng; *vegfaa* - 0.5, 1, 2, 4 and 8 ng; control morpholino - 0.5, 1, 2, 8 and 16 ng. We also injected the buffer alone to serve as a blank control. Each sample was collected at 32 hours post-fertilization (hpf). 5 embryos were pooled into one biological replicate, and a total of 4 biological replicates was obtained for each sample. We extracted RNA with TRIzol^®^ (Ambion). The aqueous phase was passed through a RNA purification column (Zymogen) and treated with RQ1 DNase (Promega). Samples were re-extracted by acid phenol:chloroform (Ambion) followed by chloroform back-extraction. Biological replicates with poor RNA quality or insufficient RNA amount were discarded. Nevertheless, each group had at least 3 biological replicates. 150 ng of RNA from each sample was converted to cDNA by High Capacity RNA-to-cDNA Kit (Applied Biosystems). Primer sequences and optimized cycling conditions for *rpl13*, *gapdh*, *isg20*, *isg15*, *casp8*, *phlda3* and *tp53Δ113* can be found in Table S1.

### Statistical Analyses of qPCR assays

All statistical analyses were performed with R statistical software. All C_T_ values were normalized to *gapdh* and compared to uninjected samples (-ΔΔC_T_). Morpholino dose is transformed to a log-scale (base 2). A dose-response curve test was applied using the *LL.4* function from drc package in R to calculate the upper limit, lower limit and the slope.

Otherwise, linear plots were tested using linear regression to estimate the slope. For plotting of graphs, both the in-built plot function of R as well as ggplot2 package [25] were utilized.

### Whole mount *in situ* hybridization (WISH)

We injected 0.5 ng of *egfl7* or control morpholino into 1-cell WT embryos (ABs). Embryos were collected at 32 hpf and fixed with 4% PFA in 1X PBST (1X PBS, 0.1% Tween-20), followed by standard WISH protocol [26]. Primers used to clone *isg20* for synthesizing the *in situ* probe are: 5’-GGAGAATCATGGGAAGTGGA-3’; and 5’-GGCATTGAGGTTGGCAGTAT-3’.

## Results

### Integration of Affymetrix microarray gene expression datasets

We retrieved 63 microarray gene expression datasets from Gene Expression Omnibus (GEO) by keyword search. The majority of microarrays used were manufactured by Agilent (n = 34) or Affymetrix (n = 19). Duplicated datasets, datasets with no accompanying publications and datasets in incorrect file formats (Agilent, n = 12; Affymetrix, n= 8) were removed from further analyses. We applied algorithms to eliminate batch effects and other technical noise in order to integrate datasets generated using the Affymetrix microarray platform (n = 11) into one large dataset. Principal component analysis (PCA) of this dataset revealed that developmental stage and tissue source were the main factors driving the clustering in the principal components (PC) 1 and 2, respectively (Fig 1A). In addition, we observed an occasional segregation of morphants (MO) from controls (CO or WT) for a given cluster group (Fig 1B).

**Fig 1.**
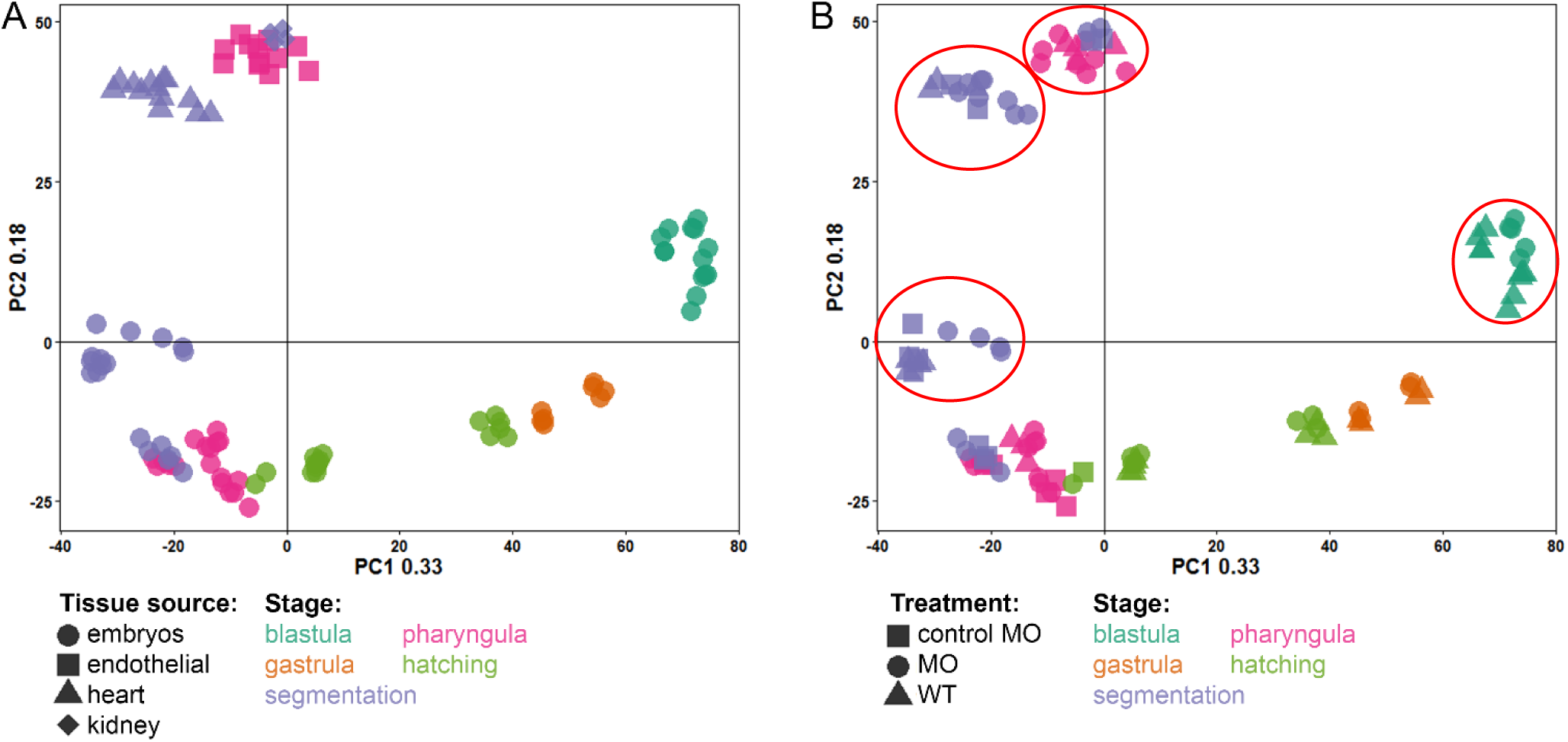
Principal component analysis (PCA) of integrated gene expression data compiled from morpholino studies in zebrafish. (A) Gene expression patterns are clustered by developmental stage along PC1 and clustered by tissue source along PC2. (B) Some morpholino experiments show morphants clustering marginally away from controls (demarcated in red).

### Feature selection uncovers activation of ISGs and the p53 signaling pathway

To identify genes differentially regulated between uninjected controls or buffer injected controls (WT) and morphants, we fed the entire dataset into a supervised machine learning algorithm. The model assigns a higher score to a gene if it performs better at predicting morphants. Among the various models of feature selection, we found that the linear feature selection model tended to perform best (Table S2). We then tested the accuracy of the model and short-listed the optimal number of genes that successfully distinguish the morphants from controls (see Materials and Methods). We refer to this set of genes as “top performing genes” (Table S3).

We detected *tp53* as one of these top performing genes upregulated in morphants compared to controls, corroborating previous findings [9]. Surprisingly, we found another gene, *isg20*, which scored higher than *tp53*, raising the possibility that interferon signaling was also activated in morphants. Furthermore, we found biological processes such as apoptosis and various cell stress responses enriched when further analyzing the top performing genes (Fig 2).

**Fig 2.**
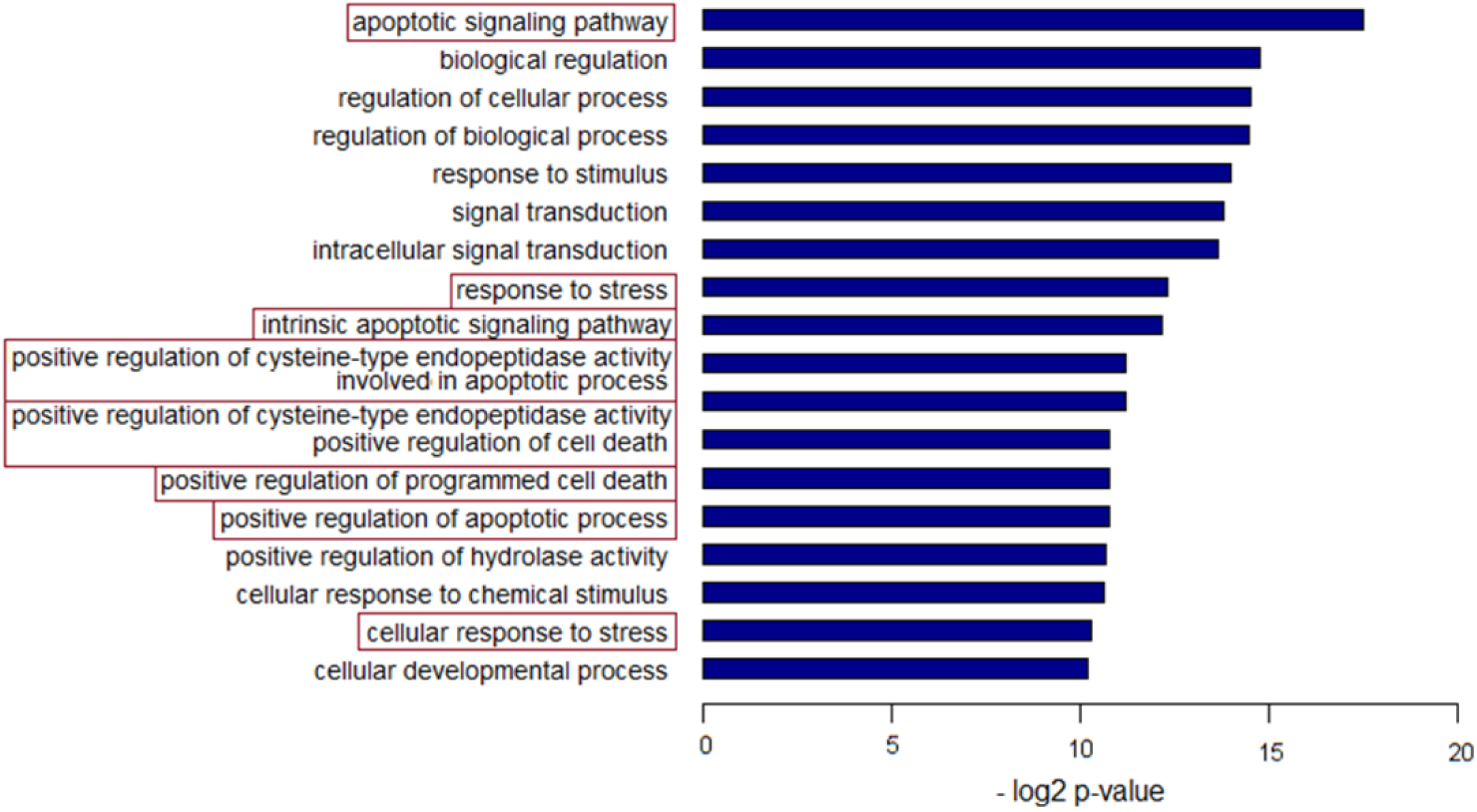
Gene set enrichment analysis. Biological processes such as apoptosis and the p53 signaling pathway are enriched in the top performing genes (S3 Table).

### ISG and p53 signaling activation in morphants are most pronounced at the segmentation stage

As mentioned earlier, the samples can be separated by developmental stage along the first PC, as expected by the dynamic nature of gene expression in the developing zebrafish embryo [27–28]. Thus, we proceeded to analyze our dataset by stratifying based on developmental stage. Using a two-way ANOVA approach, we identified 157 genes that were significantly (α = 0.001) differentially regulated between morphants and controls, depending on developmental stage (S4 Table). Interestingly, we found that *isg15*, *isg20*, *casp8*, *mdm2*, *gadd45aa*, *phlda3* and *tp53* were profoundly upregulated during segmentation stages in morphants compared to controls (Fig 3). Interestingly, Robu and colleagues similarly reported the activation of *isg20l2*, *phlda3* and *casp8* in their microarray assay (Fig 9C in [9]). Genes that are differentially regulated at other developmental stages can be found in the File S1 – Section 3.

**Fig 3.**
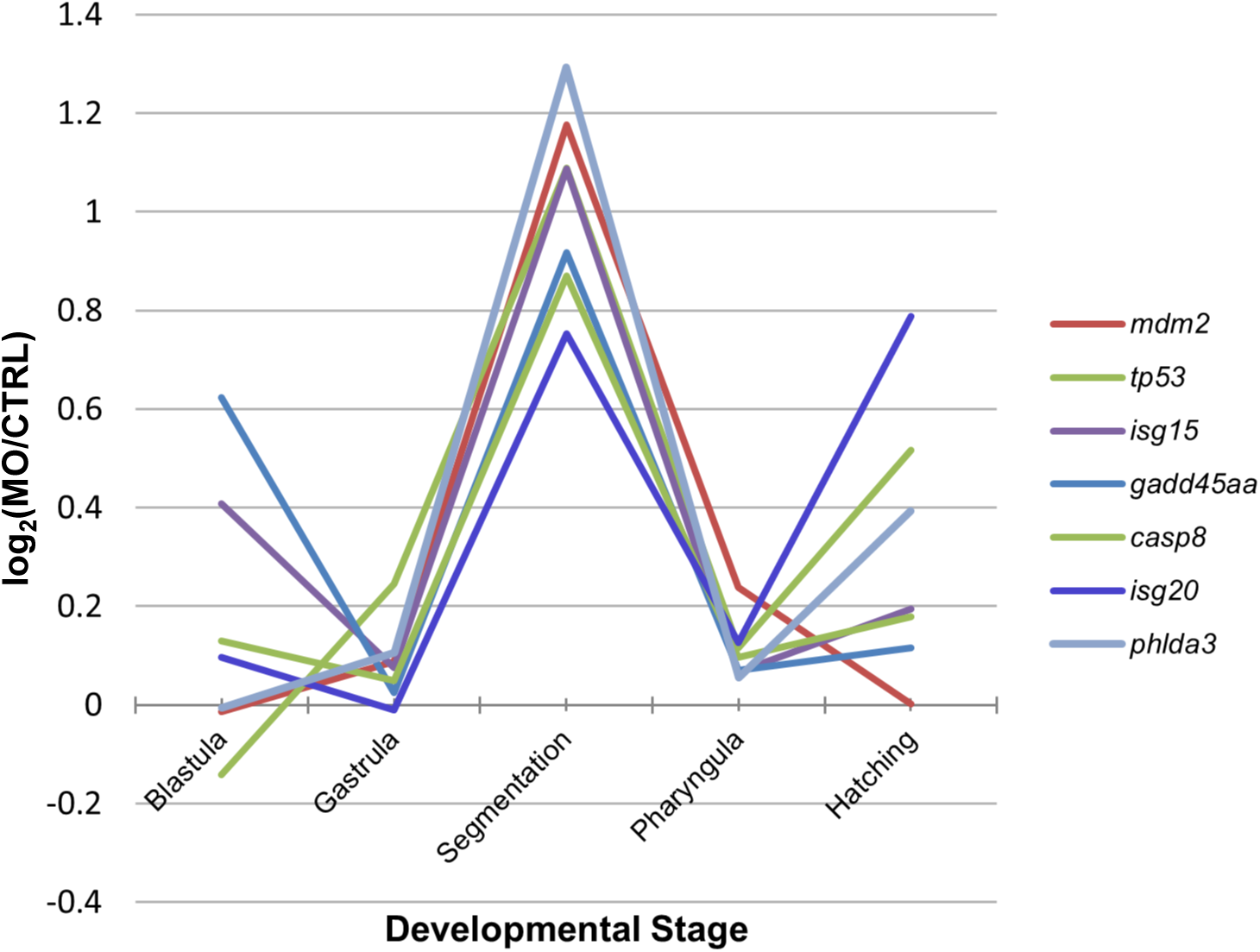
Relative fold change of gene expression plotted against developmental stage. Genes from the p53 signaling pathway (*tp53*, *mdm2*, *gadd45aa*, *phlda3*), the caspase cascade (*casp8*), as well as interferon stimulated genes (*isg15*, *isg20*) are upregulated in morphants at the segmentation stage. Morphants (MO) were compared to uninjected, buffer-injected or control morpholino (CTRL), depending on the design of the experiment from the microarray datasets.

### Analysis of Agilent microarray datasets confirms the activation of ISGs and the p53 signaling pathway in morphants

The Agilent microarray datasets were composed of different platform configurations which makes data integration challenging and could potentially introduce artefacts into the post-processed data. Therefore, we individually normalized each microarray dataset to investigate the expression levels of *isg15*, *isg20*, *casp8*, *mdm2*, *gadd45aa* and *tp53*. Indeed, these genes are similarly upregulated in some morphants compared to control embryos, and many of these morphants were studied at segmentation and pharyngula stages (Fig 4).

**Fig 4.**
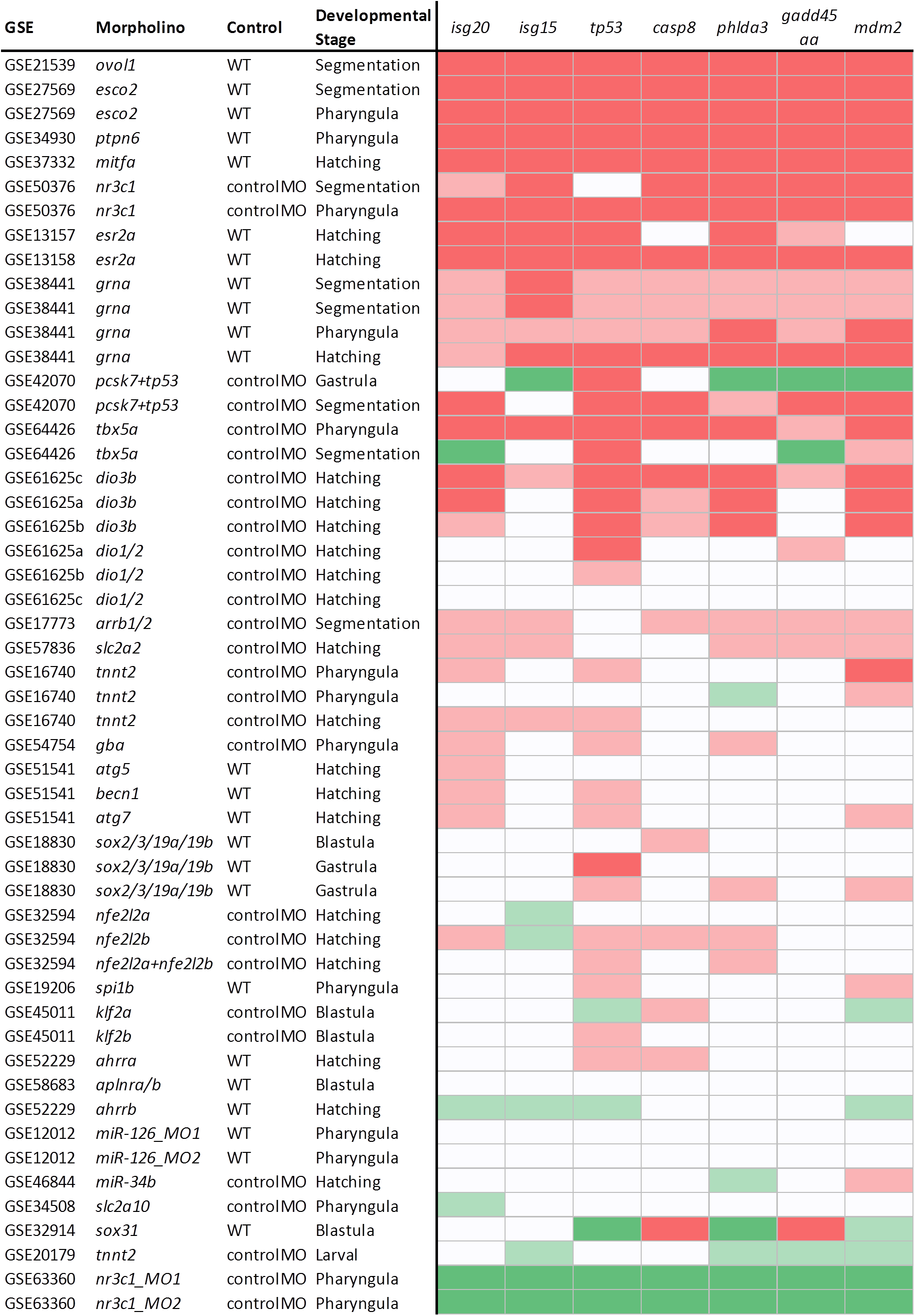
Analysis of *isg20, isg15, tp53, casp8, phlda3, gadd45aa* and *mdm2* expression from Agilent microarray datasets. Differential expression of *isg20, isg15, tp53, casp8, phlda3, gadd45aa* and *mdm2* between morphants and controls. Red indicates significant upregulation (*P* < 0.05); green indicates significant downregulation (*P* < 0.05). The darker shade of red/green indicates logFC > 1.

### ISGs are upregulated by increasing doses of some morpholinos

We injected increasing doses of *egfl7*, *vegfaa* and standard control morpholinos, followed by qPCR assays of *isg20, isg15, tp53Δ113* [9]*, phlda3* and *casp8*. We found that these genes were upregulated by increasing doses of *egfl7* morpholino, and in some assayed genes, a sigmoid curve was observed (Fig 5). In contrast, these genes were not upregulated by increasing doses of *vegfaa* or standard control morpholinos (Fig 5). However, when we injected 8 or 16 ng of standard control morpholino, not only did the injected embryos exhibit upregulation of the aforementioned genes (Fig 5), but these embryos also exhibited a delay in angiogenesis (Fig S1). We performed whole-mount *in situ* hybridization to investigate the *isg20* expression pattern in *egfl7* morphants, and found that *isg20* was upregulated throughout the embryo (Fig S2), but not restricted to the vascular endothelium which expresses *egfl7* specifically [23].

**Fig 5.**
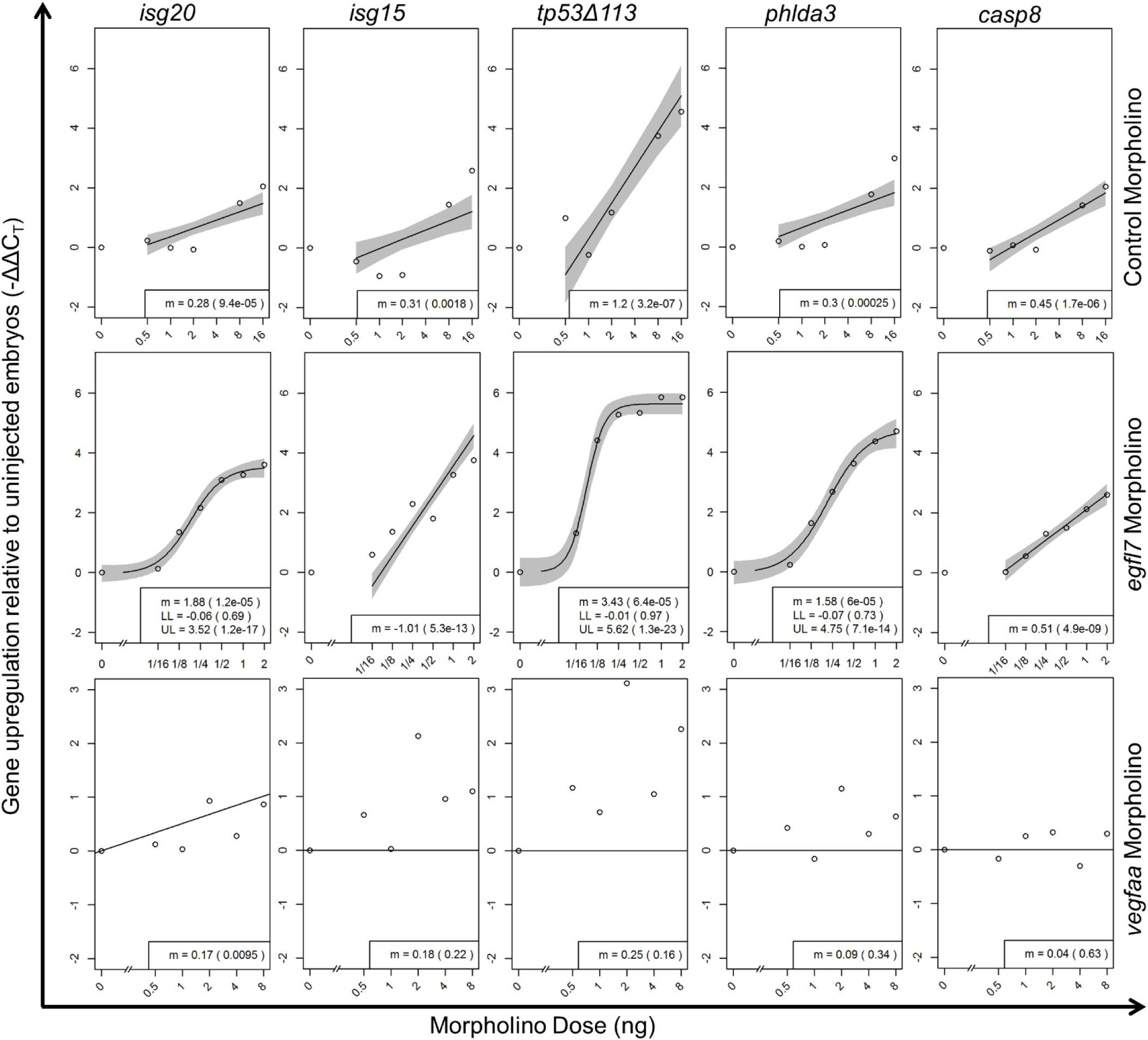
ISGs and stress response genes are upregulated in a morpholino dose-dependent manner. qPCR assays for *isg20, isg15, tp53Δ113, phlda3* and *casp8* expression in 32 hpf zebrafish embryos injected with increasing doses of standard control morpholino, *egfl7* morpholino and *vegfaa* morpholino. N = 4 per morpholino per dose. m - slope; LL - lower limit of sigmoid curve; UL - upper limit of sigmoid curve. *P*-values for individual parameters in parantheses. Grey shaded region reflects the 95% confidence band of the curve.

## Discussion

Morpholinos are widely used as a tool to inactivate gene and non-coding RNA function. However, we do not yet fully understand the responses of a cell to morpholinos. Indeed, several reported side-effects of morpholinos, including *tp53Δ113* upregulation [9], mis-splicing of transcripts and induction of the innate immune response [10], could complicate conclusions arising from morpholino studies. Similarly, our current study identified genes related to the innate immune response which are upregulated across multiple zebrafish morphants as well as zebrafish embryos injected with an excess of standard control morpholino.

Apart from the p53 signaling pathway [9], we further identified the activation of ISGs. *isg20* was previously shown to be induced by the presence of dsRNA [29–30]. dsRNA can be recognized by receptors including Toll-Like Receptor 3 (TLR3), RIG1 and MDA5, which trigger downstream interferon type-I (IFN-I) signaling [31–33]. Interestingly, we detected activation of the IL-8 signaling pathway in our top upregulated genes (data not shown), suggesting the activation of TLRs and IFN-I, possibly by an RNA-morpholino heteroduplex.

Large-scale gene expression analyses have been used to identify gene expression signatures that correlate to tumor progression, and with thorough validation, have become diagnostic markers and/or drug targets [34–35]. Moreover, public databases, such as Oncomine [36], have been established to mine out markers across multiple studies. We applied a similar strategy to multiple gene expression datasets from zebrafish morpholino studies in order to identify gene expression signatures that are common to morpholino-injected zebrafish, and found that many genes in the innate immunity pathway are upregulated in morphants.

Additional work will be needed to understand the underlying biology of these genes and how they can affect morpholino experiments. Clearer understanding of morpholino effects, should help scientists make informed decisions when conducting morpholino experiments (see also latest guidelines in ref. 37), and to facilitate interpretation of data arising from these experiments.

## Supporting information

## Acknowledgements

We would like to thank Sven Reischauer, Oliver Stone, and Hans-Martin Maischein for their insights regarding morpholinos, Mario Looso and his team for feedback and suggestions on bioinformatics, Pedro Moura for insightful discussions regarding dsRNA induction of IFN-I, and Andrea Rossi, Zacharias Kontarakis and Albert Wang for comments on the manuscript.

## Supporting Information

**Fig S1. 36 hpf zebrafish embryo trunks.** Micrographs of zebrafish trunks marked by an endothelial transgenic marker, and overlays on the bright field channel. Embryos injected with 8 ng control morpholino exhibit a mild delay in intersegmental vessel sprouting, and this delay is more pronounced in embryos injected with 16 ng control morpholino.

**Fig S2. Whole mount *in situ* for *isg20* expression in *egfl7* morphants and controls**.

**Table S1. Primers for qPCR assays of *isg20, isg15, tp53Δ113, casp8, phlda3* and *gapdh* expression levels.**

**Table S2. Prediction accuracy of various machine learning methods**.

**Table S3. Top performing genes ranked by t-test statistics and linear models**.

**Table S4. Genes that are significantly differentially regulated in a two-way ANOVA analysis**.

**File S1. Supplementary Information**.

