## Supplementary material for "Induction of Interferon-Stimulated Genes and Cellular Stress Pathways by Morpholinos"

***TABLE OF CONTENTS***

PAGE

1. **Data collection and annotation**
   1.1 Platform selection and integration	3
   1.2 GEO repository details and experiment information from literature	4
   1.3 Affymetrix microarray profiles used for integrated data analysis	6
   1.4 Agilent microarray profiles used for integrated data analysis	8
2. **Data processing and analysis of Affymetrix GPL1319 datasets**
   2.1 Data normalization and integration of Affymetrix GPL1319	11
   2.2 Recursive feature elimination algorithm	15
   2.3 Hierarchical clustering - Affymetrix	16
   2.4 Optimized parameters of embedded feature selectors	18
   2.5 Recursive feature elimination - Affymetrix	19
   2.6 Differentially expressed genes in Gastrula and Hatching stages	20
   2.7 *tp53* probes in Affymetrix microarray	26
3. **Data processing and meta-analysis of Agilent datasets**
   3.1 Agilent datasets used for meta-analysis	27
   3.2 Probe IDs mapping to genes used in meta-analysis	28
   3.3 *tp53* probes in Agilent microarrays	29

1 **Data Collection and annotation**
  1.1 Platform selection and integration

**Table S1.1.1 –**Affymetrix and Agilent platforms from GEO repository. Each GPL is shown with the number of corresponding morpholino studies correctly associated with a microarray profile. Highlighted the platforms selected for the study

| **GPL*** | **Number of studies** | **Technology** |
| --- | --- | --- |
| 1319 | 11 | Affymetrix |
| 14664 | 4 | Agilent |
| 6457 | 4 | Agilent |
| 7735 | 5 | Agilent custom array |
| 7302 | 3 | Agilent |
| 6563 | 1 | Agilent |
| 7301 | 2 | Agilent |
| 10042 | 1 | Agilent |
| 13390 | 2 | Agilent custom array |
| 15180 | 1 | Agilent custom array |
| 15450 | 1 | Agilent custom array |
| 18725 | 1 | Agilent |

*GEO platform accession

1.2 GEO repository details and experiment information from literature

**Table S1.2.1 –** Microarray data curation: the attributes below are assigned to each microarray dataset. GPL, GSE, GSM, Title and Pubmed attributes are used as identification for individual studies obtained from GEO repository. MO target, MOseq, COseq, MOtarget, source, Hpf, Treatment, MO type, MO dose and phenotype are extracted from the annexed scientific article. Dev. Stage and MO dose are created additionally from the provided information.

| Attribute | Type | Levels | Description |
| --- | --- | --- | --- |
| GPL | Descriptive | - | Type of technology |
| GSE | Descriptive | - | Study ID (coupled to unique publication) |
| GSM | Descriptive | - | Data set ID |
| Title | Descriptive | - | Data set description |
| Pubmed | Descriptive | - | Associated publication ID |
| MO target | Descriptive | - | Gene targeted by morpholino |
| MOseq | Descriptive | - | Morpholino sequence |
| COseq | Descriptive | - | Sequence of the control morpholino (if available) |
| MO target | Descriptive | - | Gene targeted by morpholino |
| Source | Categorical | kidney, endothelial cells, hearts, whole embryo, embryo trunk | Tissue type |
| Hpf | Continuous | - | Age (hours post fertilization) of the sampled embryos |
| Treatment | Categorical | Morpholino, Control | Type of treatment |
| MO type | Categorical | Translation, splicing | Mechanism of gene product inhibition |
| MO dose | Continuous | - | Nanograms of morpholino injected |
| Phenotype | Categorical | Normal, various | Phenotype induced by the injection |
| Dev. Stage | Categorical | Blastula, Gastrula, Segmentation, Pharyngula, Hatching | Developmental stage of the sampled embryos* |
| MO dose pM | Continuous | - | PicoMolar dose of the injected morpholino** |

* From: Zebrafish developmental staging series: <http://zfin.org/zf_info/zfbook/stages/> - ZFin data base

Detailed description of the zebrafish developmental stages in section 2.3

** Calculated from the morpholino sequence molecular weight and the injected quantity (MO dose). Molecular weights:

Morpholino ring: 87.12 g/mol, A subunit residue: 339.3 g/mole, C subunit residue: 315.3 g/mole, G subunit residue: 355.3 g/mole, T subunit residue: 330.3 g/mole, Additional weight of the ends of the chain:	101 g/mole

1.3 Affymetrix microarray profiles used for integrated data analysis

**Table S1.3.1** – Relevant information used for the integrated microarray study – Affymetrix

| GSE | GSM | Description | Treatment* | | Source | MO dose pM | Dev. stage |
| --- | --- | --- | --- | --- | --- | --- | --- |
| GSE12012 | GSM303576 | zebrafish_48hpf_control_replicate1 | WT | endothelial | | 0 | Pharyngula |
| GSE12012 | GSM303577 | zebrafish_48hpf_miR-126 MO-1_replicate1 | miR-126 | endothelial | | 0.455 | Pharyngula |
| GSE12012 | GSM303578 | zebrafish_48hpf_miR-126 MO-2_replicate1 | miR-126 | endothelial | | 0.563 | Pharyngula |
| GSE12012 | GSM303579 | zebrafish_48hpf_control_replicate2 | WT | endothelial | | 0 | Pharyngula |
| GSE12012 | GSM303580 | zebrafish_48hpf_miR-126 MO-1_replicate2 | miR-126 | endothelial | | 0.455 | Pharyngula |
| GSE12012 | GSM303581 | zebrafish_48hpf_miR-126 MO-2_replicate2 | miR-126 | endothelial | | 0.563 | Pharyngula |
| GSE12012 | GSM303582 | zebrafish_48hpf_control_replicate3 | WT | endothelial | | 0 | Pharyngula |
| GSE12012 | GSM303583 | zebrafish_48hpf_miR-126 MO-1_replicate3 | miR-126 | endothelial | | 0.455 | Pharyngula |
| GSE12012 | GSM303584 | zebrafish_48hpf_miR-126 MO-2_replicate3 | miR-126 | endothelial | | 0.563 | Pharyngula |
| GSE12012 | GSM303585 | zebrafish_48hpf_control_replicate4 | WT | endothelial | | 0 | Pharyngula |
| GSE12012 | GSM303586 | zebrafish_48hpf_miR-126 MO-1_replicate4 | miR-126 | endothelial | | 0.455 | Pharyngula |
| GSE12012 | GSM303587 | zebrafish_48hpf_miR-126 MO-2_replicate4 | miR-126 | endothelial | | 0.563 | Pharyngula |
| GSE13157 | GSM329381 | Embryos injected with 15uM ERbeta2 MO biological repl. 1 | esr2a | embryos | | 0.015 | Hatching |
| GSE13157 | GSM329382 | Embryos injected with 15uM ERbeta2 MO biological repl. 2 | esr2a | embryos | | 0.015 | Hatching |
| GSE13157 | GSM329383 | Embryos injected with 15uM coMO biological repl. 1 | CO | embryos | | 0.015 | Hatching |
| GSE13157 | GSM329384 | Embryos injected with 15uM coMO biological repl. 2 | CO | embryos | | 0.015 | Hatching |
| GSE13157 | GSM329385 | Embryos uninjected biological repl. 1 (15uM) | WT | embryos | | 0 | Hatching |
| GSE13157 | GSM329386 | Embryos uninjected biological repl. 2 (15uM) | WT | embryos | | 0 | Hatching |
| GSE13158 | GSM329399 | Embryos injected with 50uM ERbeta2 MO biological repl. 1 | esr2a | embryos | | 0.05 | Hatching |
| GSE13158 | GSM329400 | Embryos injected with 50uM ERbeta2 MO biological repl. 2 | esr2a | embryos | | 0.05 | Hatching |
| GSE13158 | GSM329401 | Embryos injected with 50uM coMO biological repl. 1 | CO | embryos | | 0.05 | Hatching |
| GSE13158 | GSM329402 | Embryos injected with 50uM coMO biological repl. 2 | CO | embryos | | 0.05 | Hatching |
| GSE13158 | GSM329403 | Embryos uninjected biological repl. 1 (50uM) | WT | embryos | | 0 | Hatching |
| GSE13158 | GSM329404 | Embryos uninjected biological repl. 2 (50uM) | WT | embryos | | 0 | Hatching |
| GSE16740 | GSM419445 | RNA from embryos of 3 groups of Wildtype AB zebrafish at 36 hpf_1 | CO | embryos | | 0.38** | Pharyngula |
| GSE16740 | GSM419446 | RNA from embryos of 3 groups of Wildtype AB zebrafish at 36 hpf_2 | CO | embryos | | 0.38** | Pharyngula |
| GSE16740 | GSM419447 | RNA from embryos of 3 groups of Wildtype AB zebrafish at 36 hpf_3 | CO | embryos | | 0.38** | Pharyngula |
| GSE16740 | GSM419448 | RNA from embryos of 3 groups of tnnt2 morpholino fish at 36 hpf_1 | tnnt2 | embryos | | 0.377** | Pharyngula |
| GSE16740 | GSM419449 | RNA from embryos of 3 groups of tnnt2 morpholino fish at 36 hpf_2 | tnnt2 | embryos | | 0.377** | Pharyngula |
| GSE16740 | GSM419450 | RNA from embryos of 3 groups of tnnt2 morpholino fish at 36 hpf_3 | tnnt2 | embryos | | 0.377** | Pharyngula |
| GSE16740 | GSM419451 | RNA from embryos of 3 groups of Wildtype AB zebrafish at 48 hpf_1 | CO | embryos | | 0.38** | Pharyngula |
| GSE16740 | GSM419452 | RNA from embryos of 3 groups of Wildtype AB zebrafish at 48 hpf_2 | CO | embryos | | 0.38** | Pharyngula |
| GSE16740 | GSM419453 | RNA from embryos of 3 groups of Wildtype AB zebrafish at 48 hpf_3 | CO | embryos | | 0.38** | Pharyngula |
| GSE16740 | GSM419454 | RNA from embryos of 3 groups of tnnt2 morpholino fish at 48 hpf_1 | tnnt2 | embryos | | 0.377** | Pharyngula |
| GSE16740 | GSM419455 | RNA from embryos of 3 groups of tnnt2 morpholino fish at 48 hpf_2 | tnnt2 | embryos | | 0.377** | Pharyngula |
| GSE16740 | GSM419456 | RNA from embryos of 3 groups of tnnt2 morpholino fish at 48 hpf_3 | tnnt2 | embryos | | 0.377** | Pharyngula |
| GSE16740 | GSM419457 | RNA from embryos of 3 groups of Wildtype AB zebrafish at 60 hpf_1 | CO | embryos | | 0.38** | Hatching |
| GSE16740 | GSM419458 | RNA from embryos of 3 groups of Wildtype AB zebrafish at 60 hpf_2 | CO | embryos | | 0.38** | Hatching |
| GSE16740 | GSM419459 | RNA from embryos of 3 groups of Wildtype AB zebrafish at 60 hpf_3 | CO | embryos | | 0.38** | Hatching |
| GSE16740 | GSM419460 | RNA from embryos of 3 groups of tnnt2 morpholino fish at 60 hpf_1 | tnnt2 | embryos | | 0.377** | Hatching |
| GSE16740 | GSM419461 | RNA from embryos of 3 groups of tnnt2 morpholino fish at 60 hpf_2 | tnnt2 | embryos | | 0.377** | Hatching |
| GSE16740 | GSM419462 | RNA from embryos of 3 groups of tnnt2 morpholino fish at 60 hpf_3 | tnnt2 | embryos | | 0.377** | Hatching |
| GSE18830 | GSM466790 | Zebrafish wt embryo_ 30%E_biological rep1 | WT | embryos | | 0 | Blastula |
| GSE18830 | GSM466791 | Zebrafish wt embryo_ 30%E_biological rep2 | WT | embryos | | 0 | Blastula |
| GSE18830 | GSM466792 | Zebrafish wt embryo_ 75%E_biological rep1 | WT | embryos | | 0 | Gastrula |
| GSE18830 | GSM466793 | Zebrafish wt embryo_ 75%E_biological rep2 | WT | embryos | | 0 | Gastrula |
| GSE18830 | GSM466794 | Zebrafish wt embryo_ TB_biological rep1 | WT | embryos | | 0 | Gastrula |
| GSE18830 | GSM466795 | Zebrafish wt embryo_ TB_biological rep2 | WT | embryos | | 0 | Gastrula |
| GSE18830 | GSM466796 | Zebrafish QKD embryo_ 30%E_biological rep1 | sox2/3/19a/19b | embryos | | 0.068 | Blastula |
| GSE18830 | GSM466797 | Zebrafish QKD embryo_ 30%E_biological rep2 | sox2/3/19a/19b | embryos | | 0.068 | Blastula |
| GSE18830 | GSM466798 | Zebrafish QKD embryo_ 75%E_biological rep1 | sox2/3/19a/19b | embryos | | 0.068 | Gastrula |
| GSE18830 | GSM466799 | Zebrafish QKD embryo_ 75%E_biological rep2 | sox2/3/19a/19b | embryos | | 0.068 | Gastrula |
| GSE18830 | GSM466800 | Zebrafish QKD embryo_TB_biological rep1 | sox2/3/19a/19b | embryos | | 0.068 | Gastrula |
| GSE18830 | GSM466801 | Zebrafish QKD embryo_TB_biological rep2 | sox2/3/19a/19b | embryos | | 0.068 | Gastrula |
| GSE21539 | GSM537962 | zebrafish 12hpf control-1 | WT | embryos | | 0 | Segmentation |
| GSE21539 | GSM537963 | zebrafish 12hpf control-2 | WT | embryos | | 0 | Segmentation |
| GSE21539 | GSM537964 | zebrafish 12hpf control-3 | WT | embryos | | 0 | Segmentation |
| GSE21539 | GSM537965 | zebrafish 12hpf Ovo1 morphant-1 | Ovo1 | embryos | | 0.286 | Segmentation |
| GSE21539 | GSM537966 | zebrafish 12hpf Ovo1 morphant-2 | Ovo1 | embryos | | 0.286 | Segmentation |
| GSE21539 | GSM537967 | zebrafish 12hpf Ovo1 morphant-3 | Ovo1 | embryos | | 0.286 | Segmentation |
| GSE27569 | GSM683599 | control embryos at 24hpf, biological rep 1 | WT | embryos | | 0 | Segmentation |
| GSE27569 | GSM683600 | control embryos at 24hpf, biological rep 2 | WT | embryos | | 0 | Segmentation |
| GSE27569 | GSM683601 | control embryos at 24hpf, biological rep 3 | WT | embryos | | 0 | Segmentation |
| GSE27569 | GSM683602 | control embryos at 24hpf, biological rep 4 | WT | embryos | | 0 | Segmentation |
| GSE27569 | GSM683603 | esco2 MO injected embryos at 24hpf, biological rep 1 | esco2 | embryos | | 0.189 | Segmentation |
| GSE27569 | GSM683604 | esco2 MO injected embryos at 24hpf, biological rep 2 | esco2 | embryos | | 0.189 | Segmentation |
| GSE27569 | GSM683605 | esco2 MO injected embryos at 24hpf, biological rep 3 | esco2 | embryos | | 0.189 | Segmentation |
| GSE27569 | GSM683606 | esco2 MO injected embryos at 24hpf, biological rep 4 | esco2 | embryos | | 0.189 | Segmentation |
| GSE27569 | GSM683607 | control embryos at 48hpf, biological rep 1 | WT | embryos | | 0 | Pharyngula |
| GSE27569 | GSM683608 | control embryos at 48hpf, biological rep 3 | WT | embryos | | 0 | Pharyngula |
| GSE27569 | GSM683609 | control embryos at 48hpf, biological rep 4 | WT | embryos | | 0 | Pharyngula |
| GSE27569 | GSM683610 | esco2 MO injected embryos at 48hpf, biological rep 1 | esco2 | embryos | | 0.189 | Pharyngula |
| GSE27569 | GSM683611 | esco2 MO injected embryos at 48hpf, biological rep 2 | esco2 | embryos | | 0.189 | Pharyngula |
| GSE27569 | GSM683612 | esco2 MO injected embryos at 48hpf, biological rep 3 | esco2 | embryos | | 0.189 | Pharyngula |
| GSE27569 | GSM683613 | esco2 MO injected embryos at 48hpf, biological rep 4 | esco2 | embryos | | 0.189 | Pharyngula |
| GSE32914 | GSM814795 | zebrafish_SB_4.3h_1 | Sox31 | embryos | | 1.2 | Blastula |
| GSE32914 | GSM814796 | zebrafish_SB_4.3h_2 | Sox31 | embryos | | 1.2 | Blastula |
| GSE32914 | GSM814797 | zebrafish_SB_4.3h_3 | Sox31 | embryos | | 1.2 | Blastula |
| GSE32914 | GSM814798 | zebrafish_WT_4.3h_1 | WT | embryos | | 0 | Blastula |
| GSE32914 | GSM814799 | zebrafish_WT_4.3h_2 | WT | embryos | | 0 | Blastula |
| GSE32914 | GSM814800 | zebrafish_WT_4.3h_3 | WT | embryos | | 0 | Blastula |
| GSE32914 | GSM814801 | zebrafish_WT_2.5h_1 | WT | embryos | | 0 | Blastula |
| GSE32914 | GSM814802 | zebrafish_WT_2.5h_2 | WT | embryos | | 0 | Blastula |
| GSE32914 | GSM814803 | zebrafish_WT_4h_1 | WT | embryos | | 0 | Blastula |
| GSE32914 | GSM814804 | zebrafish_WT_4h_2 | WT | embryos | | 0 | Blastula |
| GSE46844 | GSM1139092 | Multiciliated cell_control_rep1 | CO | kidney | | 1 | Hatching |
| GSE46844 | GSM1139093 | Multiciliated cell_control_rep2 | CO | kidney | | 1 | Hatching |
| GSE46844 | GSM1139094 | Multiciliated cell_miR-34B morphants_rep1 | mir-34B | kidney | | 1 | Hatching |
| GSE46844 | GSM1139095 | Multiciliated cell_miR-34B morphants_rep2 | mir-34B | kidney | | 1 | Hatching |
| GSE51541 | GSM1247632 | Uninjected control biological replicate 1 | WT | heart | | 0 | Hatching |
| GSE51541 | GSM1247633 | Uninjected control biological replicate 2 | WT | heart | | 0 | Hatching |
| GSE51541 | GSM1247634 | Uninjected control biological replicate 3 | WT | heart | | 0 | Hatching |
| GSE51541 | GSM1247635 | Control Morpholino biological replicate 2 | CO | heart | | 0.65 | Hatching |
| GSE51541 | GSM1247636 | Control Mopholino biological replicate 3 | CO | heart | | 0.65 | Hatching |
| GSE51541 | GSM1247637 | atg5 Mopholino biological replicate 1 | atg5 | heart | | 0.65 | Hatching |
| GSE51541 | GSM1247638 | atg5 Mopholino biological replicate 2 | atg5 | heart | | 0.65 | Hatching |
| GSE51541 | GSM1247639 | atg5 Mopholino biological replicate 3 | atg5 | heart | | 0.65 | Hatching |
| GSE51541 | GSM1247640 | becn1 Mopholino biological replicate 1 | becn1 | heart | | 0.65 | Hatching |
| GSE51541 | GSM1247641 | becn1 Mopholino biological replicate 2 | becn1 | heart | | 0.65 | Hatching |
| GSE51541 | GSM1247642 | becn1 Mopholino biological replicate 3 | becn1 | heart | | 0.65 | Hatching |
| GSE51541 | GSM1247643 | atg7 Mopholino biological replicate 1 | atg7 | heart | | 0.65 | Hatching |
| GSE51541 | GSM1247644 | atg7 Mopholino biological replicate 2 | atg7 | heart | | 0.65 | Hatching |
| GSE8800 | GSM218665 | MO | C1q-like | embryos | | 0.473 | Segmentation |
| GSE8800 | GSM218666 | Cont MO | CO | embryos | | 0.474 | Segmentation |

* Treatment

*Control:* **WT** – uninjected, **CO** – injected morpholino sequence

*Morpholino treatments:* **microRNA**- mir34B, mir126, **genes:** C1q-like, atg7, becn1, atg5, Sox31, esco2, Ovo1, sox2/3/19a/19b, tnnt2, esr2a

**Dose information extracted from Sehnert et al., 2002

**Table S1.3.2** – Number of microarrays per attribute factor. The table is a summary of Table S1.3

| Developmental stage | Blastula: 14  Gastrula: 8  Segmentation: 16  Pharyngula: 31  Hatching: 35 |
| --- | --- |
| Treatment | Morpholino: 52  Control: 52 (WT:34 -CO:18) |
| Source | Embryos: 75  Kidney: 4  Hearts: 13  Endothelial cells: 12 |
| Dosage: | 0-0.189pM: 56  0.289-0.65pM: 41  1-1.2pM: 7 |

1.4 Agilent microarray profiles used for integrated data analysis

**Table S1.4.1** – Relevant information used for the integrated microarray study- Agilent

| GSE | GSM | Description | Treatment* | | Source | MO dose pM | Dev. stage |
| --- | --- | --- | --- | --- | --- | --- | --- |
| GSE32594 | GSM807955 | Injection of control MO at 1-4 cell stage. DMSO exposure at 48 hpf. RNA sampled at 52 hpf. | CO | embryos | | 0.05 | Hatching |
| GSE32594 | GSM807956 | Injection of control MO at 1-4 cell stage. DMSO exposure at 48 hpf. RNA sampled at 52 hpf. | CO | embryos | | 0.05 | Hatching |
| GSE32594 | GSM807957 | Injection of control MO at 1-4 cell stage. DMSO exposure at 48 hpf. RNA sampled at 52 hpf. | CO | embryos | | 0.05 | Hatching |
| GSE32594 | GSM807961 | Injection of Nrf2a MO at 1-4 cell stage. DMSO exposure at 48 hpf. RNA sampled at 52 hpf. | Nrf2a | embryos | | 0.025 | Hatching |
| GSE32594 | GSM807962 | Injection of Nrf2a MO at 1-4 cell stage. DMSO exposure at 48 hpf. RNA sampled at 52 hpf. | Nrf2a | embryos | | 0.025 | Hatching |
| GSE32594 | GSM807963 | Injection of Nrf2a MO at 1-4 cell stage. DMSO exposure at 48 hpf. RNA sampled at 52 hpf. | Nrf2a | embryos | | 0.025 | Hatching |
| GSE32594 | GSM807967 | Injection of Nrf2a and Nrf2b MO at 1-4 cell stage. DMSO exposure at 48 hpf. RNA sampled at 52 hpf. | Nrf2a/Nrf2b | embryos | | 0.05 | Hatching |
| GSE32594 | GSM807968 | Injection of Nrf2a and Nrf2b MO at 1-4 cell stage. DMSO exposure at 48 hpf. RNA sampled at 52 hpf. | Nrf2a/Nrf2b | embryos | | 0.05 | Hatching |
| GSE32594 | GSM807969 | Injection of Nrf2a and Nrf2b MO at 1-4 cell stage. DMSO exposure at 48 hpf. RNA sampled at 52 hpf. | Nrf2a/Nrf2b | embryos | | 0.05 | Hatching |
| GSE32594 | GSM807973 | Injection of Nrf2b MO at 1-4 cell stage. DMSO exposure at 48 hpf. RNA sampled at 52 hpf. | Nrf2b | embryos | | 0.025 | Hatching |
| GSE32594 | GSM807974 | Injection of Nrf2b MO at 1-4 cell stage. DMSO exposure at 48 hpf. RNA sampled at 52 hpf. | Nrf2b | embryos | | 0.025 | Hatching |
| GSE32594 | GSM807975 | Injection of Nrf2b MO at 1-4 cell stage. DMSO exposure at 48 hpf. RNA sampled at 52 hpf. | Nrf2b | embryos | | 0.025 | Hatching |
| GSE42070 | GSM1031965 | whole fish, 6h, control, replicate 1 | CO_R | embryos | | 0.5 | Gastrula |
| GSE42070 | GSM1031966 | whole fish, 6h, control, replicate 2 | CO_R | embryos | | 0.5 | Gastrula |
| GSE42070 | GSM1031967 | whole fish, 6h, control, replicate 3 | CO_R | embryos | | 0.5 | Gastrula |
| GSE42070 | GSM1031968 | whole fish, 24h, control, replicate 1 | CO_R | embryos | | 0.5 | Segmentation |
| GSE42070 | GSM1031969 | whole fish, 24h, control, replicate 2 | CO_R | embryos | | 0.5 | Segmentation |
| GSE42070 | GSM1031970 | whole fish, 24h, control, replicate 3 | CO_R | embryos | | 0.5 | Segmentation |
| GSE42070 | GSM1031971 | whole fish, 6h, PCSK7+P53 morpholinos, replicate 1 | PCSK7/P53 | embryos | | 1.25 | Gastrula |
| GSE42070 | GSM1031972 | whole fish, 6h, PCSK7+P53 morpholinos, replicate 2 | PCSK7/P53 | embryos | | 1.25 | Gastrula |
| GSE42070 | GSM1031973 | whole fish, 6h, PCSK7+P53 morpholinos, replicate 3 | PCSK7/P53 | embryos | | 1.25 | Gastrula |
| GSE42070 | GSM1031974 | whole fish, 24h, PCSK7+P53 morpholinos, replicate 1 | PCSK7/P53 | embryos | | 1.25 | Segmentation |
| GSE42070 | GSM1031975 | whole fish, 24h, PCSK7+P53 morpholinos, replicate 2 | PCSK7/P53 | embryos | | 1.25 | Segmentation |
| GSE42070 | GSM1031976 | whole fish, 24h, PCSK7+P53 morpholinos, replicate 3 | PCSK7/P53 | embryos | | 1.25 | Segmentation |
| GSE45012 | GSM1095807 | FoxD5 Morpholino injection | FoxD5 | embryos | | 0.38 | Blastula |
| GSE45012 | GSM1095808 | FoxD5 Morpholino injection | FoxD5 | embryos | | 0.38 | Blastula |
| GSE45012 | GSM1095813 | FoxD5 Morpholino injection | FoxD5 | embryos | | 0.38 | Blastula |
| GSE20179 | GSM506241 | Whole embryo TNNT2sp morphants | tnnt2 | embryos | | 0.471 | Hatching |
| GSE20179 | GSM506241 | Whole embryo TNNT2sp morphants | CO | embryos | | 0.471 | Hatching |
| GSE20179 | GSM506242 | Whole embryo TNNT2sp morphants | tnnt2 | embryos | | 0.471 | Hatching |
| GSE20179 | GSM506242 | Whole embryo TNNT2sp morphants | CO | embryos | | 0.471 | Hatching |
| GSE20179 | GSM506243 | Whole embryo TNNT2sp morphants | tnnt2 | embryos | | 0.471 | Hatching |
| GSE20179 | GSM506243 | Whole embryo TNNT2sp morphants | CO | embryos | | 0.471 | Hatching |
| GSE20179 | GSM506244 | Whole embryo TNNT2sp morphants | tnnt2 | embryos | | 0.471 | Hatching |
| GSE20179 | GSM506244 | Whole embryo TNNT2sp morphants | CO | embryos | | 0.471 | Hatching |
| GSE24934 | GSM612962 | MO2-ers2a embryos at 8 hours post fertilization | CO | embryos | | 0.974 | Gastrula |
| GSE24934 | GSM612962 | MO2-ers2a embryos at 8 hours post fertilization | Ers2a | embryos | | 0.974 | Gastrula |
| GSE24934 | GSM612963 | MO2-ers2a embryos at 48 hours post fertilization | CO | embryos | | 0.974 | Pharyngula |
| GSE24934 | GSM612963 | MO2-ers2a embryos at 48 hours post fertilization | Ers2a | embryos | | 0.974 | Pharyngula |
| GSE25517 | GSM627464 | MO2-nr3c1-5m, 5hpf | CO | embryos | | 0.783 | Blastula |
| GSE25517 | GSM627464 | MO2-nr3c1-5m, 5hpf | MO | embryos | | 0.783 | Blastula |
| GSE25517 | GSM627465 | WT, 5hpf | WT | embryos | | 0.783 | Blastula |
| GSE25517 | GSM627465 | WT, 5hpf | MO | embryos | | 0.783 | Blastula |
| GSE25517 | GSM627466 | MO2-nr3c1-5m, 10hpf | CO | embryos | | 0.783 | Gastrula |
| GSE25517 | GSM627466 | MO2-nr3c1-5m, 10hpf | MO | embryos | | 0.783 | Gastrula |
| GSE25517 | GSM627467 | WT, 10hpf | WT | embryos | | 0.783 | Gastrula |
| GSE25517 | GSM627467 | WT, 10hpf | MO | embryos | | 0.783 | Gastrula |
| GSE38441 | GSM942059 | 16 hpf trunk region of zebrafish embryos: WT vs MO-grnA injected Replicate 1 | WT | trunk | | 0.023 | Segmentation |
| GSE38441 | GSM942059 | 16 hpf trunk region of zebrafish embryos: WT vs MO-grnA injected Replicate 1 | MO | trunk | | 0.023 | Segmentation |
| GSE38441 | GSM942060 | 16 hpf trunk region of zebrafish embryos: WT vs MO-grnA injected Replicate 2 | WT | trunk | | 0.023 | Segmentation |
| GSE38441 | GSM942060 | 16 hpf trunk region of zebrafish embryos: WT vs MO-grnA injected Replicate 2 | MO | trunk | | 0.023 | Segmentation |
| GSE38441 | GSM942061 | 16 hpf trunk region of zebrafish embryos: WT vs MO-grnA injected Replicate 3 | WT | trunk | | 0.023 | Segmentation |
| GSE38441 | GSM942061 | 16 hpf trunk region of zebrafish embryos: WT vs MO-grnA injected Replicate 3 | MO | trunk | | 0.023 | Segmentation |
| GSE38441 | GSM942062 | 24 hpf trunk region of zebrafish embryos: WT vs MO-grnA injected Replicate 1 | WT | trunk | | 0.023 | Segmentation |
| GSE38441 | GSM942062 | 24 hpf trunk region of zebrafish embryos: WT vs MO-grnA injected Replicate 1 | MO | trunk | | 0.023 | Segmentation |
| GSE38441 | GSM942063 | 24 hpf trunk region of zebrafish embryos: WT vs MO-grnA injected Replicate 2 | WT | trunk | | 0.023 | Segmentation |
| GSE38441 | GSM942063 | 24 hpf trunk region of zebrafish embryos: WT vs MO-grnA injected Replicate 2 | MO | trunk | | 0.023 | Segmentation |
| GSE38441 | GSM942064 | 24 hpf trunk region of zebrafish embryos: WT vs MO-grnA injected Replicate 3 | WT | trunk | | 0.023 | Segmentation |
| GSE38441 | GSM942064 | 24 hpf trunk region of zebrafish embryos: WT vs MO-grnA injected Replicate 3 | MO | trunk | | 0.023 | Segmentation |
| GSE38441 | GSM942065 | 48 hpf trunk region of zebrafish embryos: WT vs MO-grnA injected Replicate 1 | WT | trunk | | 0.023 | Pharyngula |
| GSE38441 | GSM942065 | 48 hpf trunk region of zebrafish embryos: WT vs MO-grnA injected Replicate 1 | MO | trunk | | 0.023 | Pharyngula |
| GSE38441 | GSM942066 | 48 hpf trunk region of zebrafish embryos: WT vs MO-grnA injected Replicate 2 | WT | trunk | | 0.023 | Pharyngula |
| GSE38441 | GSM942066 | 48 hpf trunk region of zebrafish embryos: WT vs MO-grnA injected Replicate 2 | MO | trunk | | 0.023 | Pharyngula |
| GSE38441 | GSM942067 | 48 hpf trunk region of zebrafish embryos: WT vs MO-grnA injected Replicate 3 | WT | trunk | | 0.023 | Pharyngula |
| GSE38441 | GSM942067 | 48 hpf trunk region of zebrafish embryos: WT vs MO-grnA injected Replicate 3 | MO | trunk | | 0.023 | Pharyngula |
| GSE38441 | GSM942068 | 72 hpf trunk region of zebrafish embryos: WT vs MO-grnA injected Replicate 1 | WT | trunk | | 0.023 | Hatching |
| GSE38441 | GSM942068 | 72 hpf trunk region of zebrafish embryos: WT vs MO-grnA injected Replicate 1 | MO | trunk | | 0.023 | Hatching |
| GSE38441 | GSM942069 | 72 hpf trunk region of zebrafish embryos: WT vs MO-grnA injected Replicate 2 | WT | trunk | | 0.023 | Hatching |
| GSE38441 | GSM942069 | 72 hpf trunk region of zebrafish embryos: WT vs MO-grnA injected Replicate 2 | MO | trunk | | 0.023 | Hatching |
| GSE38441 | GSM942070 | 72 hpf trunk region of zebrafish embryos: WT vs MO-grnA injected Replicate 3 | WT | trunk | | 0.023 | Hatching |
| GSE38441 | GSM942070 | 72 hpf trunk region of zebrafish embryos: WT vs MO-grnA injected Replicate 3 | MO | trunk | | 0.023 | Hatching |
| GSE45011 | GSM1095795 | Klf2a Morpholino injection | Klf2a | embryos | | 0.38 | Blastula |
| GSE45011 | GSM1095799 | Klf2a+Klf2b Morpholino injection | Klf2a/Klf2b | embryos | | 0.19 | Blastula |
| GSE45011 | GSM1095800 | Klf2a+Klf2b Morpholino injection | Klf2a/Klf2b | embryos | | 0.19 | Blastula |
| GSE45011 | GSM1095796 | Klf2a Morpholino injection | Klf2a | embryos | | 0.38 | Blastula |
| GSE45011 | GSM1095797 | Klf2b Morpholino injection | Klf2b | embryos | | 0.375 | Blastula |
| GSE45011 | GSM1095793 | Control Morpholino injection | CO | embryos | | 0.38 | Blastula |
| GSE45011 | GSM1095794 | Control Morpholino injection | CO | embryos | | 0.38 | Blastula |
| GSE45011 | GSM1095798 | Klf2b Morpholino injection | Klf2b | embryos | | 0.375 | Blastula |
| GSE45012 | GSM1095801 | FoxD3 Morpholino injection | FoxD3 | embryos | | 0.38 | Blastula |
| GSE45012 | GSM1095802 | FoxD3+FoxD5 Morpholino injection | FoxD3/FoxD5 | embryos | | 0.19 | Blastula |
| GSE45012 | GSM1095803 | FoxD3+FoxD5 Morpholino injection | FoxD3/FoxD5 | embryos | | 0.19 | Blastula |
| GSE45012 | GSM1095804 | FoxD3 Morpholino injection | FoxD3 | embryos | | 0.38 | Blastula |
| GSE45012 | GSM1095805 | FoxD3 Morpholino injection | FoxD3 | embryos | | 0.38 | Blastula |
| GSE45012 | GSM1095806 | FoxD3+FoxD5 Morpholino injection | FoxD3/FoxD5 | embryos | | 0.19 | Blastula |
| GSE45012 | GSM1095809 | Klf4 Morpholino injection | Klf4 | embryos | | 0.38 | Blastula |
| GSE45012 | GSM1095810 | Control Morpholino injection | CO | embryos | | 0.38 | Blastula |
| GSE45012 | GSM1095811 | Klf4 Morpholino injection | Klf4 | embryos | | 0.38 | Blastula |
| GSE45012 | GSM1095812 | Control Morpholino injection | CO | embryos | | 0.38 | Blastula |

* Treatment

*Control:* **WT** – uninjected, **CO** – injected morpholino sequence

*Morpholino treatment:* **genes:** Klf4, FoxD3/FoxD5, FoxD3, Klf2b, Klf2a/Klf2b, tnnt2, FoxD5, PCSK7/P53, Nrf2b, Nrf2a/Nrf2b, Nrf2a

**Table S1.4.2** – Number of microarrays per attribute factor. The table is a summary of Table S1.5

| Developmental stage | Blastula: 25  Gastrula: 12  Segmentation: 18  Pharyngula: 8  Hatching: 26 |
| --- | --- |
| Treatment | Morpholino: 54  Control: 32 (WT:14, CO:18) |
| Source | Embryos: 65  Trunk: 24 |
| Dosage: | 0-0.19: 43  0.38-0.783: 36  0.974-1.25: 10 |

2 **Data Processing and Analysis of Affymetrix GPL1319 Datasets**
  2.1 Data normalization and integration – Affymetrix GPL1319

**Figure S2.1.1** – Affymetrix: Boxplots of the microarray intensities. Each study is presented with different color. From the top: GSE12012, GSE13157, GSE13158, GSE16740, GSE18830, GSE21539, GSE27569, GSE32914, GSE46844, GSE51541, GSE8800

A) Raw intensities

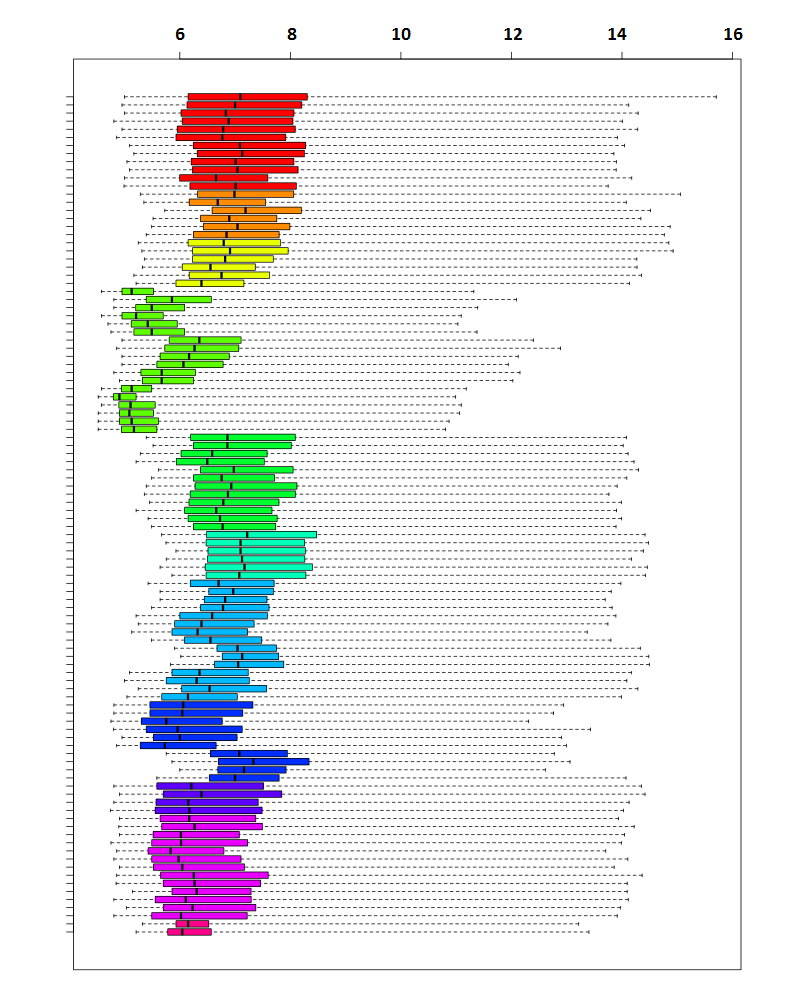

B) Intensities normalized through Single Channel Array Normalization (SCAN)

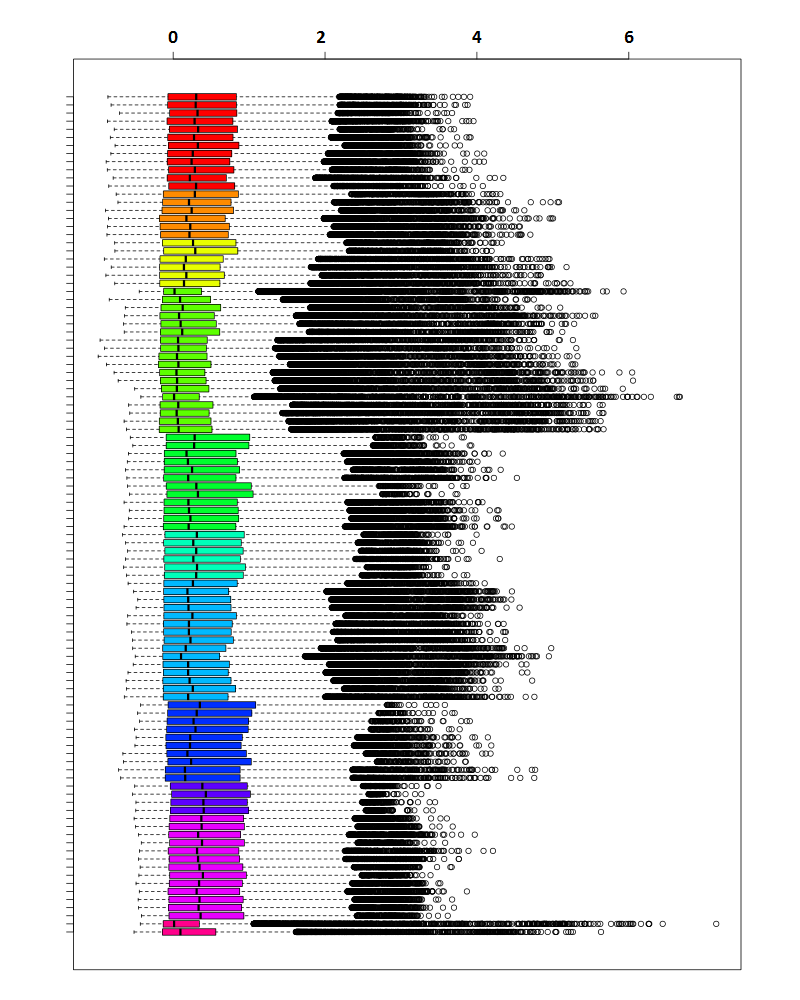

C) Intensities after Empirical Bayes “batch” standardization. The levels of the *Batch* covariate are the different studies and linear *model matrix* is included in the correction

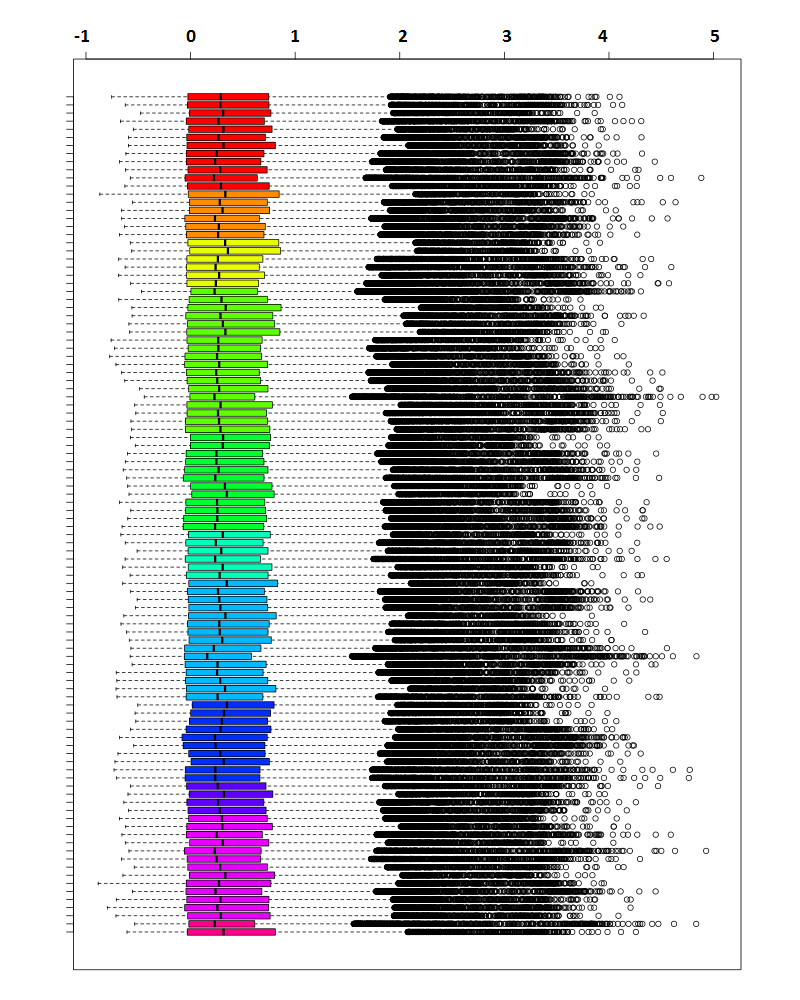

D) Q-Q plot of the intensities distribution, before and after batch correction

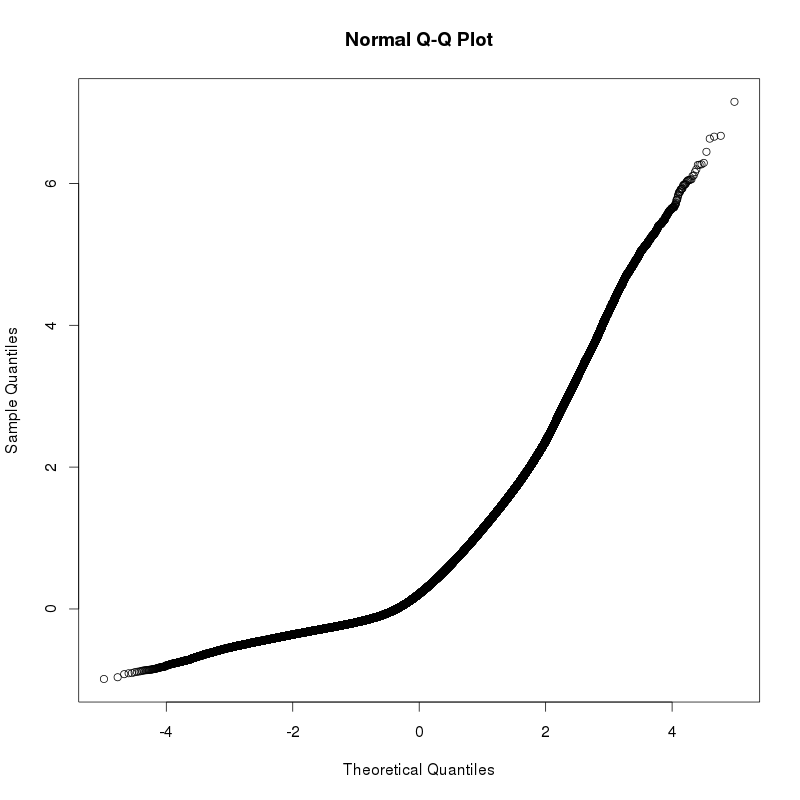

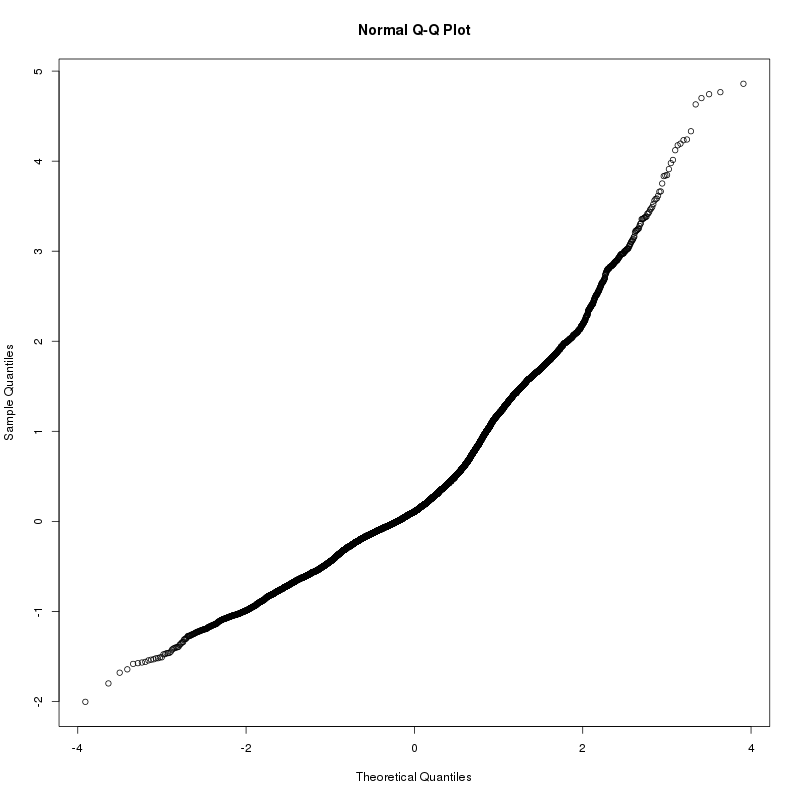

2.2 Recursive feature elimination algorithm

**Figure S2.2.1** – Schematic representation of the recursive feature elimination used for defining the optimal number of features

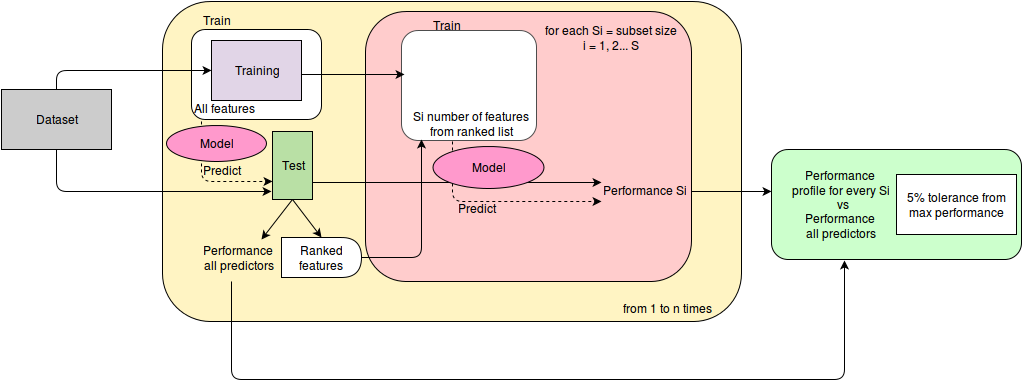

Algorithm

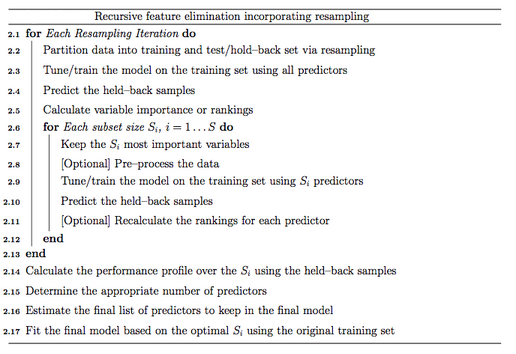

From Caret RFE documentation: <http://topepo.github.io/caret/rfe.html>

2.3 Hierarchical clustering - Affymetrix

**Figure S2.3.1** – Hierarchical clustering with Euclidian distance, Pearson correlation score, Ward´s minimum variance method (Murtagh and Legendre, 2014). The samples are labeled by treatment: Wild type, Control and Morpholino

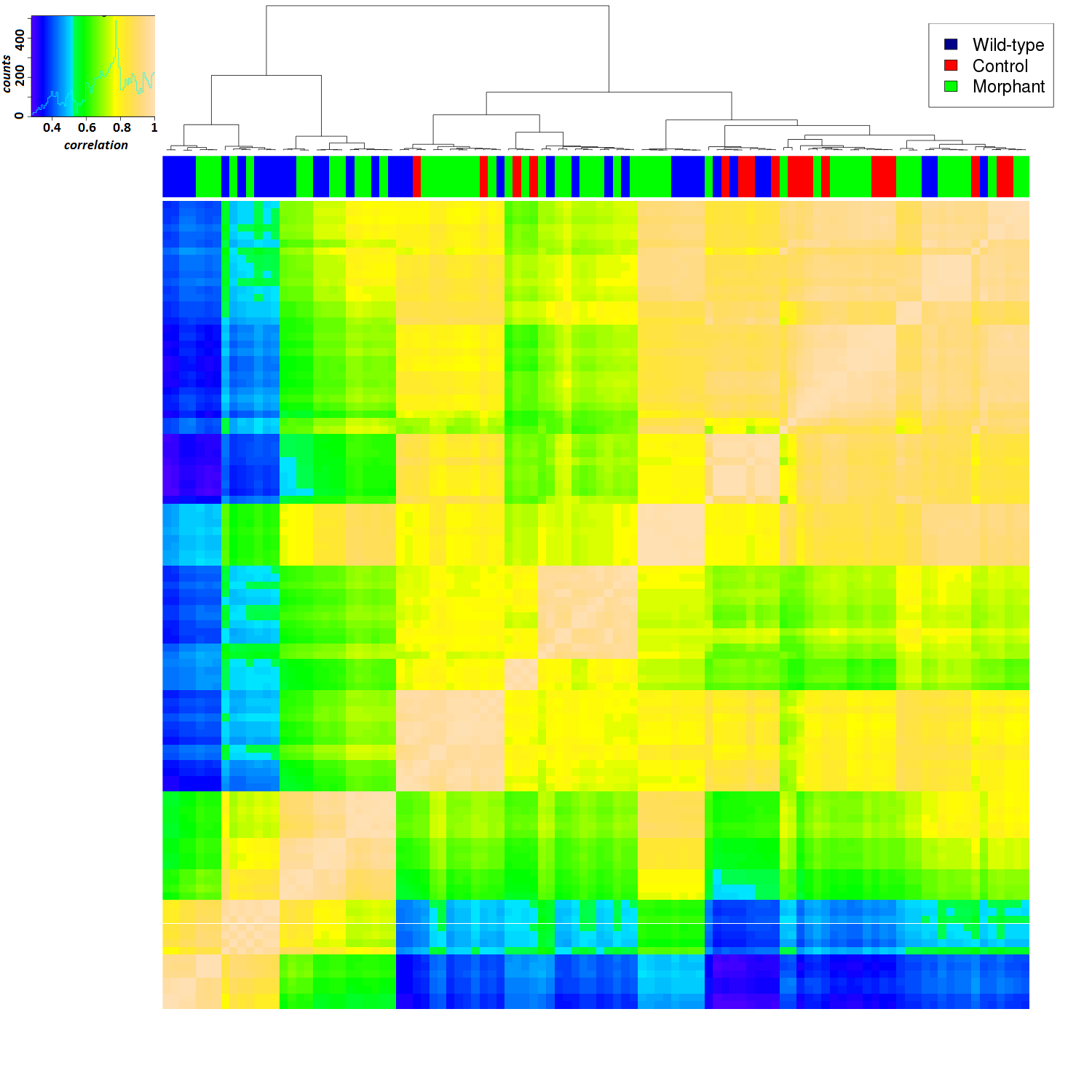

**Figure S2.3.2** – Hierarchical clustering with Euclidian distance, Pearson correlation score, Ward´s minimum variance method (Murtagh and Legendre, 2014) as the previous Figure S3.2A. The samples are labeled by developmental stage (B) or tissue type (C)

**B)**

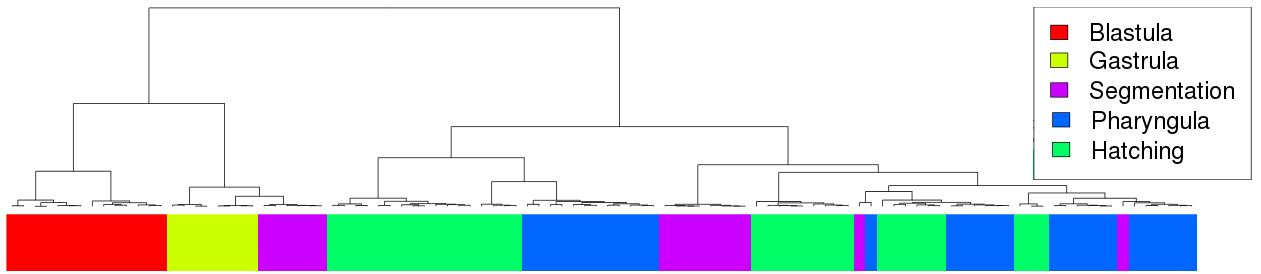

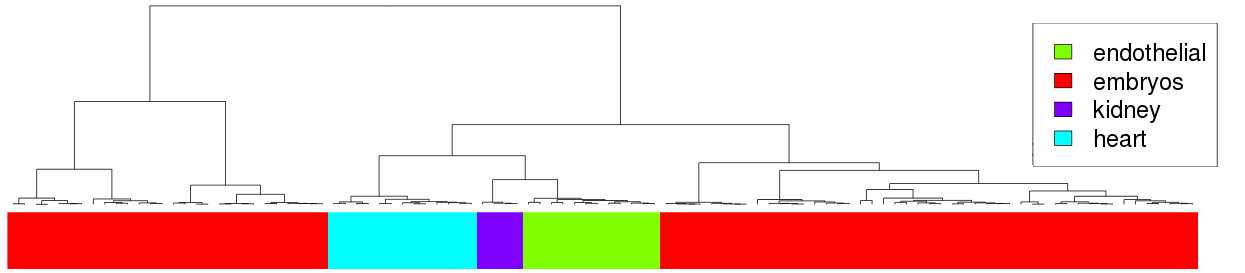
**C)**

2.4 Optimized parameters of embedded feature selectors

**Table S2.4.1** – Algorithms optimal parameters for Affymetrix

| Method | Parameters |
| --- | --- |
| GlmNet | Alpha: 0.1, Lambda: 0.1 |
| Gaussian Linear | No parameters |
| SVM Linear | C: 1 |
| Random Forest | Mtry: 3763 (max) |
| J48 | C: 0.25 |
| Naïve Bayes | fL: 0, usekerel: FALSE |
| SVM Radial kernel | Sigma: 0.00019, C: 0.5 |
| SVM Polynomial kernel | Degree: 1 , Scale: 0.01 , C: 1 |
| PAM | Threshold: 1.435 |
| KNN | K: 5 |

2.5 Recursive feature elimination – Affymetrix

**Figure S2.5.1**– **A)** Model´s accuracy for features sizes equal to 20, 30, 40, 50, 60, 100, 120, 150, 200, 250, 300 and 955. The red line shows the 5% tolerance interval **B**) Model´s accuracy for features sizes from 120 to 150. The red line shows the 5% tolerance interval

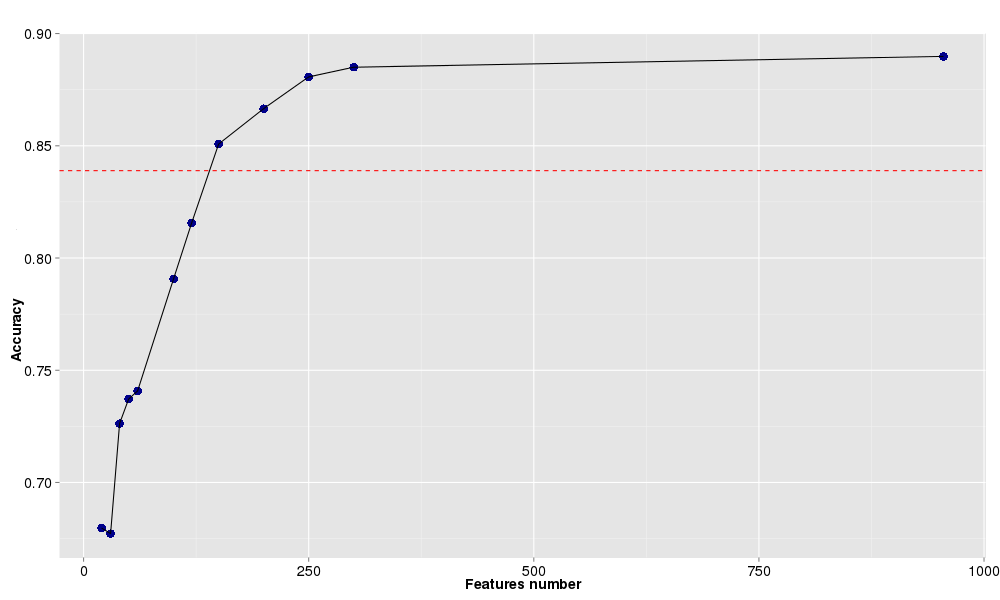

B)

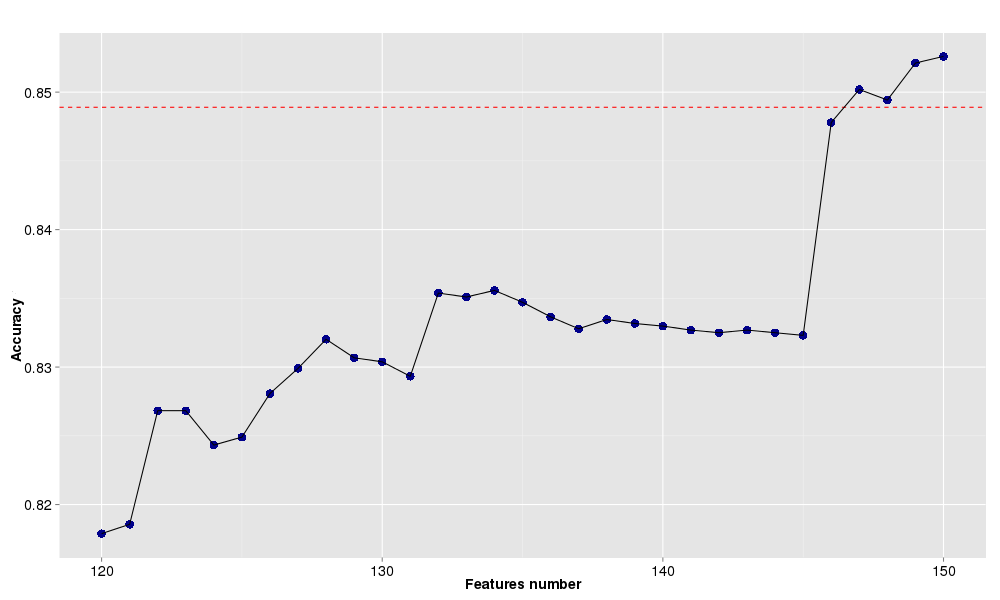

2.6 Differentially expressed genes in Gastrula and Hatching Stages

The 289 genes that were found significant in 2-WAY ANOVA were compared as Fold Change between the 2 treatments, MO and WT (log2MO- log2WT). Since the intensities are already in logarithmic scale, the fold change was calculate as the subtraction of the two values.

The values with 1.5 Fold change (0.58 in logarithmic scale) were selected as relevant change in the gene expression. Below the list of up and down regulated genes between the 2 treatments.

**Table S2.6.1A –** Genes upregulated during Gastrula stage.

| Gene Name | log2 FC | p-value |
| --- | --- | --- |
| tuba1a | 0,866 | 0,05 |
| fkbp9 | 0,752 | 0,05 |

**Table S2.6.1B –** GO annotation of genes upregulated during Gastrula stage.

| GOID | TERM | ANNOTATED_GENES |
| --- | --- | --- |
| GO:0006457 | protein folding | fkbp9 |
| GO:0008219 | cell death | tuba1a |
| GO:0022607 | cellular component assembly | tuba1a |
| GO:0065003 | macromolecular complex assembly | tuba1a |
| GO:0048646 | anatomical structure formation involved in morphogenesis | tuba1a |
| GO:0048856 | anatomical structure development | tuba1a |
| GO:0006461 | protein complex assembly | tuba1a |

**Table S2.6.2A –** Genes downregulated during Gastrula stage.

| Gene Name | log2 FC | P-value |
| --- | --- | --- |
| kctd15a | -1,263 | 0,05 |
| crabp2b | -1,246 | 0,001 |
| dlx3b | -0,966 | 0,001 |
| msxb | -0,823 | 0,05 |
| msxe | -0,749 | 0,001 |
| her9 | -0,662 | 0,05 |
| inaa | -0,611 | 0,05 |
| lmo4a | -0,605 | 0,001 |

**Table S2.6.2B –** GO annotation of genes downregulated during Gastrula stage.

| GOID | TERM | ANNOTATED_GENES |
| --- | --- | --- |
| GO:0048856 | anatomical structure development | crabp2b, dlx3b, her9, inaa, kctd15a, lmo4a, msxb, msxe |
| GO:0009058 | biosynthetic process | dlx3b, her9, kctd15a, msxb, msxe |
| GO:0034641 | cellular nitrogen compound metabolic process | dlx3b, her9, kctd15a, msxb, msxe |
| GO:0009790 | embryo development | dlx3b, kctd15a, lmo4a, msxb, msxe |
| GO:0030154 | cell differentiation | dlx3b, her9, kctd15a |
| GO:0048646 | anatomical structure formation involved in morphogenesis | dlx3b, msxb, msxe |
| GO:0007165 | signal transduction | crabp2b, kctd15a |
| GO:0006950 | response to stress | msxb |
| GO:0006810 | transport | crabp2b |
| GO:0022607 | cellular component assembly | kctd15a |
| GO:0008219 | cell death | msxb |
| GO:0065003 | macromolecular complex assembly | kctd15a |
| GO:0040007 | growth | msxb |
| GO:0006461 | protein complex assembly | kctd15a |

**Table S2.6.3A –** Genes upregulated during Hatching stage.

| Gene | log2 FC | P-value |
| --- | --- | --- |
| rrp7a | 0,812 | 0,05 |
| isg20 | 0,787 | 0,001 |
| rtcb | 0,762 | 0,05 |
| c6 | 0,756 | 0,05 |
| si:ch211-59o9.10 | 0,737 | 0,05 |
| riok3 | 0,686 | 0,05 |
| dnlz | 0,679 | 0,001 |
| ddx27 | 0,641 | 0,05 |
| psmd11a | 0,634 | 0,05 |
| TAF1C | 0,629 | 0,05 |
| snrpe | 0,628 | 0,001 |
| kctd15a | 0,618 | 0,05 |
| kctd15a | 0,618 | 0,05 |
| pak1ip1 | 0,615 | 0,05 |
| zcchc10 | 0,612 | 0,05 |
| exosc5 | 0,587 | 0,05 |

**Table S2.6.3B –** GO annotation of genes upregulated during Hatching stage.

| GOID | TERM | ANNOTATED_GENES |
| --- | --- | --- |
| GO:0034641 | cellular nitrogen compound metabolic process | ddx27, exosc5, kctd15a, rrp7a, rtcb, snrpe |
| GO:0022607 | cellular component assembly | kctd15a, psmd11a, rrp7a, snrpe |
| GO:0065003 | macromolecular complex assembly | kctd15a, psmd11a, rrp7a, snrpe |
| GO:0022618 | ribonucleoprotein complex assembly | rrp7a, snrpe |
| GO:0042254 | ribosome biogenesis | exosc5, rrp7a |
| GO:0030154 | cell differentiation | kctd15a, psmd11a |
| GO:0007165 | signal transduction | kctd15a, pak1ip1 |
| GO:0009056 | catabolic process | exosc5, psmd11a |
| GO:0006461 | protein complex assembly | kctd15a, psmd11a |
| GO:0009058 | biosynthetic process | kctd15a |
| GO:0002376 | immune system process | c6 |
| GO:0006457 | protein folding | dnlz |
| GO:0006605 | protein targeting | dnlz |
| GO:0006810 | transport | dnlz |
| GO:0006397 | mRNA processing | snrpe |
| GO:0055085 | transmembrane transport | dnlz |
| GO:0006399 | tRNA metabolic process | rtcb |
| GO:0007005 | mitochondrion organization | dnlz |
| GO:0006464 | cellular protein modification process | riok3 |
| GO:0034655 | nucleobase-containing compound catabolic process | exosc5 |
| GO:0009790 | embryo development | kctd15a |
| GO:0048856 | anatomical structure development | kctd15a |

**Table S2.6.4A –** Genes downregulated during Hatching stage.

| GeneName | log2 FC | P-value |
| --- | --- | --- |
| opn1mw1 | -2,708 | 0,001 |
| pde6h | -2,707 | 0,001 |
| prss59.2 | -2,356 | 0,001 |
| prss59.1 | -2,356 | 0,001 |
| pde6c | -2,312 | 0,001 |
| opn1sw2 | -2,231 | 0,001 |
| matn1 | -2,096 | 0,05 |
| gnat2 | -2,044 | 0,001 |
| CELA1 (1 of 7) | -1,992 | 0,001 |
| irbp | -1,88 | 0,001 |
| hsc70 | -1,581 | 0,05 |
| ctrb1 | -1,515 | 0,001 |
| try | -1,431 | 0,001 |
| zgc:136461 | -1,274 | 0,001 |
| ckmt2a | -1,245 | 0,001 |
| epd | -1,182 | 0,001 |
| ela2 | -1,112 | 0,001 |
| rcv1 | -1,111 | 0,001 |
| fabp10a | -1,105 | 0,05 |
| cpb1 | -1,1 | 0,001 |
| acta2 | -1,089 | 0,001 |
| calb2b | -1,071 | 0,05 |
| slc25a3a | -1,053 | 0,001 |
| mbpa | -0,988 | 0,001 |
| si:ch211-207l14.1 | -0,987 | 0,001 |
| elovl4b | -0,924 | 0,05 |
| tm4sf4 | -0,917 | 0,001 |
| ela3l | -0,911 | 0,001 |
| TPM1 (2 of 2) | -0,866 | 0,001 |
| guk1b | -0,808 | 0,001 |
| ugt1ab | -0,803 | 0,05 |
| rom1b | -0,799 | 0,001 |
| inaa | -0,775 | 0,05 |
| mb | -0,754 | 0,001 |
| cldn15la | -0,754 | 0,05 |
| mbpb | -0,752 | 0,05 |
| si:ch211-76l23.4 | -0,747 | 0,001 |
| cyp8b2 | -0,694 | 0,001 |
| cyp8b1 | -0,694 | 0,001 |
| si:dkey-164f24.2 | -0,677 | 0,05 |
| rpe65a | -0,669 | 0,05 |
| slc24a2 | -0,65 | 0,001 |
| myoc | -0,635 | 0,05 |
| BX957297.1 | -0,593 | 0,05 |
| eno1b | -0,584 | 0,001 |

**Table S2.6.4A –** GO annotation of genes downregulated during Hatching stage.

| GOID | TERM | ANNOTATED_GENES |
| --- | --- | --- |
| GO:0050877 | neurological system process | gnat2, opn1mw1, opn1sw2, pde6h, rom1b, rpe65a |
| GO:0048856 | anatomical structure development | acta2, inaa, matn1, mb, mbpa, pde6c |
| GO:0007165 | signal transduction | gnat2, opn1mw1, opn1sw2, pde6c, rom1b |
| GO:0044281 | small molecule metabolic process | elovl4b, eno1b, guk1b, ugt1ab |
| GO:0006810 | transport | fabp10a, mb, slc24a2, slc25a3a |
| GO:0042592 | homeostatic process | calb2b, pde6c, slc24a2 |
| GO:0006464 | cellular protein modification process | opn1mw1, opn1sw2, si:ch211-207l14.1 |
| GO:0009058 | biosynthetic process | elovl4b, ugt1ab |
| GO:0005975 | carbohydrate metabolic process | eno1b, ugt1ab |
| GO:0034641 | cellular nitrogen compound metabolic process | eno1b, guk1b |
| GO:0006629 | lipid metabolic process | cyp8b1, elovl4b |
| GO:0006950 | response to stress | mb |
| GO:0051186 | cofactor metabolic process | eno1b |
| GO:0055085 | transmembrane transport | slc24a2 |
| GO:0030154 | cell differentiation | mb |
| GO:0007267 | cell-cell signaling | slc24a2 |
| GO:0006091 | generation of precursor metabolites and energy | eno1b |
| GO:0007155 | cell adhesion | epd |
| GO:0009790 | embryo development | pde6c |

2.7 *tp53* Probes in Affymetrix Microarray

**Figure S2.6.1** – Alignment of *tp53* Affymetrix microarray probes to Zv10 *tp53* locus.

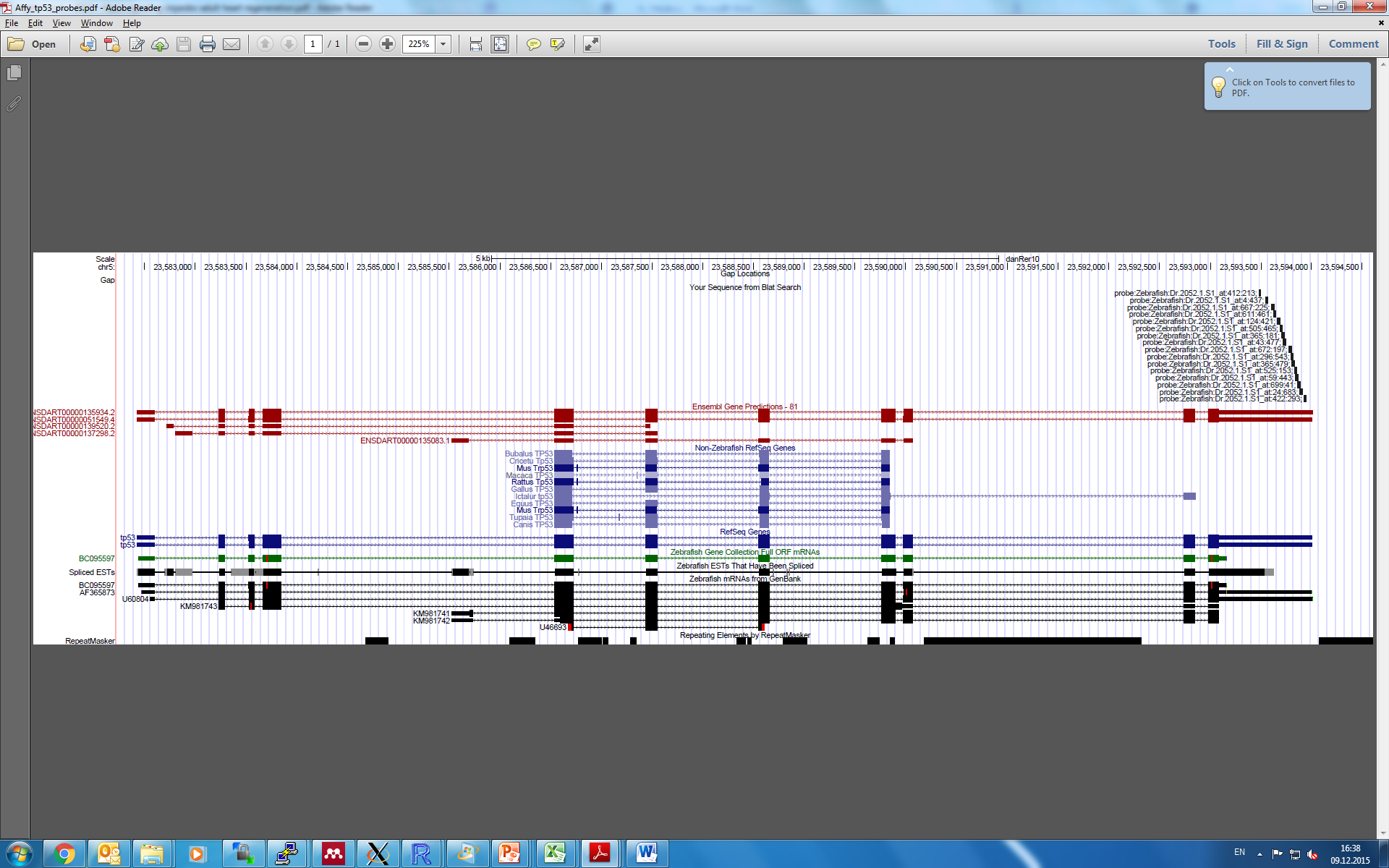

3 **Data Processing and Meta-analysis of Agilent Datasets**
  3.1 Agilent datasets used for meta-analysis

| GSE | channels | Retain? | remarks |
| --- | --- | --- | --- |
| GSE17773 | single | Yes | no replicates. pooling all MOs |
| GSE19206 | dual | Yes | some bias in MA plot |
| GSE20179 | dual | Yes | some biased A-values in 3rd array |
| GSE24934 | dual | No | wrong format. |
| GSE25517 | dual | No | some biased A-values in 2nd and 3rd arrays; WT_10hpf cannot be estimated; experiment design is too spread out. |
| GSE26708 | dual | No | wrong format |
| GSE26709 | ND | No | wrong format |
| GSE26710 | ND | No | wrong format |
| GSE32594 | single | Yes |  |
| GSE34508 | dual | Yes |  |
| GSE34930 | dual | Yes |  |
| GSE37332 | single | Yes | too few replicates. combine 48hpf and 50hpf |
| GSE38441 | dual | Yes |  |
| GSE42070 | single | Yes |  |
| GSE45011 | dual | Yes | can't estimate double MO |
| GSE45012 | dual | Yes | can't estimate double MO |
| GSE50376 | single | Yes |  |
| GSE52229 | dual | Yes |  |
| GSE54754 | single | Yes |  |
| GSE57836 | single | Yes |  |
| GSE57946 | dual | No | file error |
| GSE63360 | dual | Yes |  |

3.2 Probe IDs mapping to genes used in meta-analysis

| gene | probeID |
| --- | --- |
| *casp8* | A_15_P176121 |
| *casp8* | A_15_P115561 |
| *gadd45aa* | A_15_P171836 |
| *gadd45aa* | A_15_P113284 |
| *gadd45aa* | A_15_P116197 |
| *isg15* | A_15_P145996 |
| *isg15* | A_15_P101735 |
| *isg20* | A_15_P596582 |
| *isg20* | A_15_P108231 |
| *isg20* | A_15_P231866 |
| *tp53* | A_15_P120536 |
| *tp53* | A_15_P134831 |
| *tp53* | A_15_P102660 |
| *tp53* | A_15_P657181 |
| *tp53* | A_15_P630666 |
| *tp53* | A_15_P131466 |

3.3 *tp53* Probes in Agilent Microarrays

**Figure S3.3.1** - Alignment of *tp53* Agilent microarray probes to Zv10 *tp53* locus.

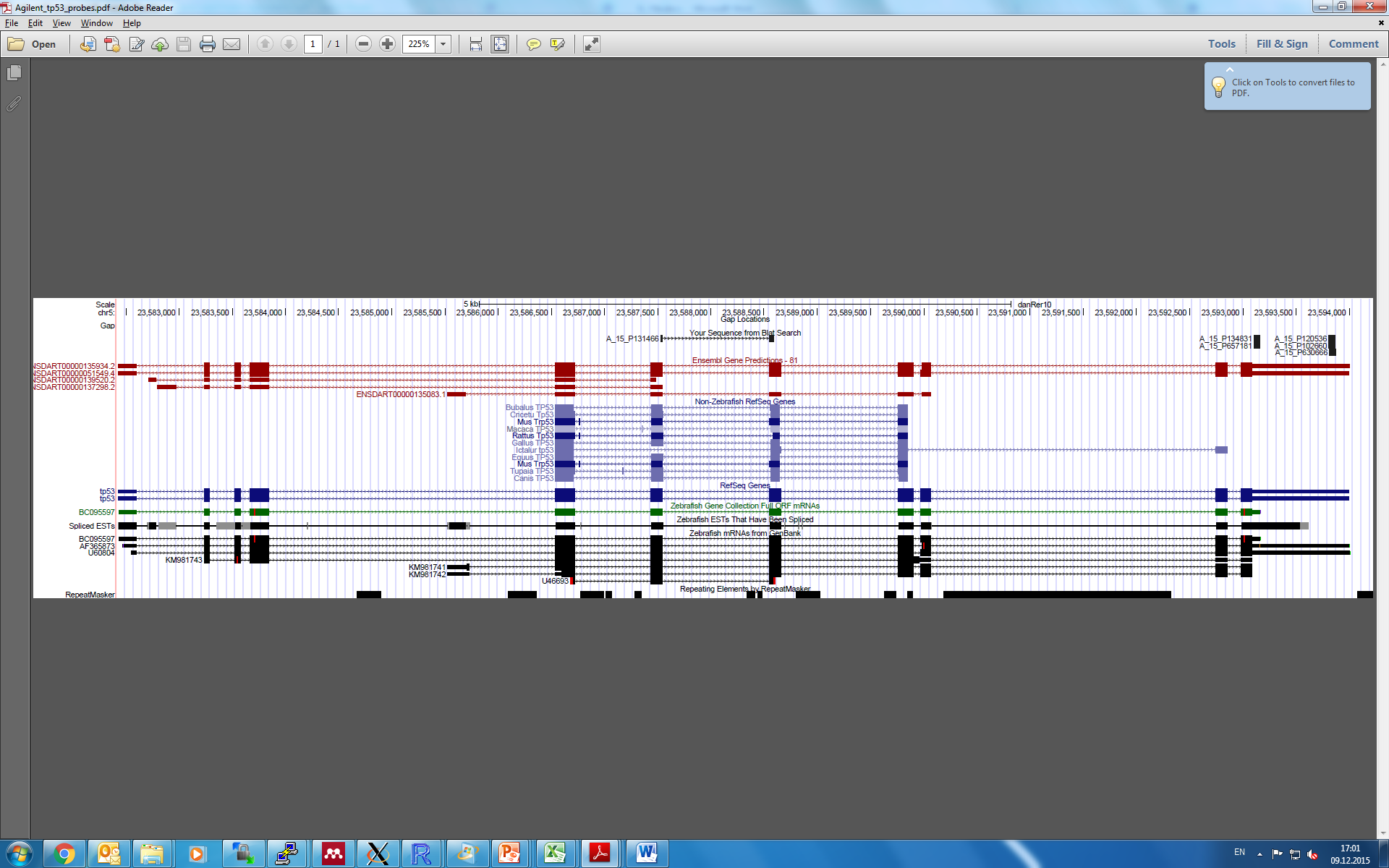
