## Supplementary figures and images for "Induction of Interferon-Stimulated Genes and Cellular Stress Pathways by Morpholinos"

### Supplementary file 6

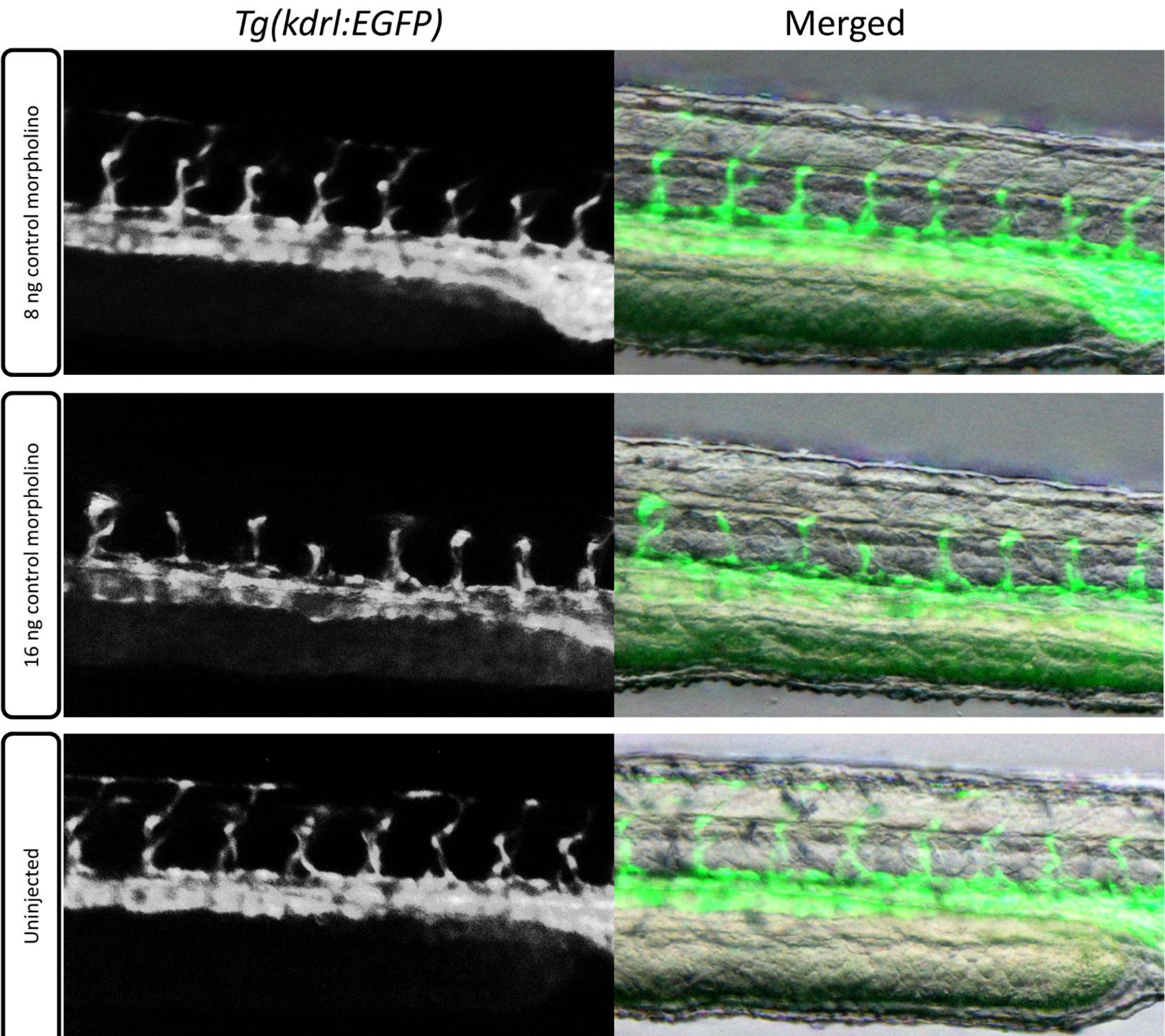

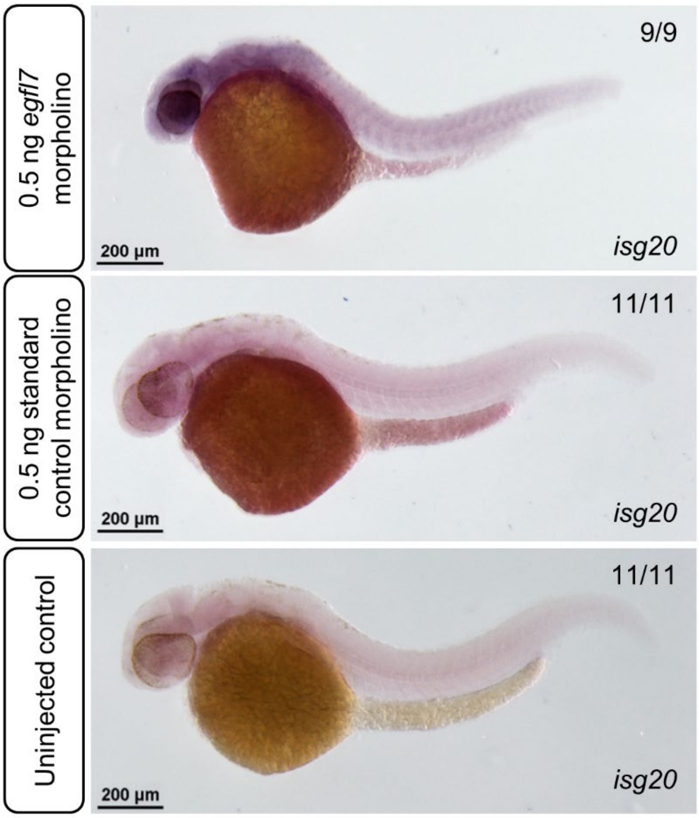
